# Regulated unbinding of ZAP70 at the T cell receptor by kinetic avidity

**DOI:** 10.1101/2020.02.12.945170

**Authors:** Jesse Goyette, David Depoil, Zhengmin Yang, Samuel A. Isaacson, Jun Allard, P. Anton van der Merwe, Katharina Gaus, Michael L Dustin, Omer Dushek

## Abstract

Protein-protein binding domains are critical in signalling networks. Src homology 2 (SH2) domains are binding domains that interact with sequences containing phosphorylated tyrosines. A subset of SH2 domain-containing proteins have tandem domains, which are thought to enhance binding affinity and specificity. However, a trade-off exists between long-lived binding and the ability to rapidly reverse signalling, which is a critical requirement of noise filtering mechanisms such as kinetic proofreading. Here, we use modelling to show that the unbinding rate of tandem, but not single, SH2 domains can be accelerated by phosphatases when tandem domains bind by a kinetic, but not a static, avidity mode. We use surface plasmon resonance to show that ZAP70, a tandem SH2 domain-containing kinase, binds kinetically to biphosphorylated peptides from the T cell antigen receptor (TCR) and that the unbinding rate can be accelerated by the phosphatase CD45. An important functional prediction of regulated unbinding is that the intracellular ZAP70/TCR half-life in T cells will be correlated to the extracellular TCR/antigen half-life and we show that this is the case in both cell lines and primary T cells. The work highlights that binding by kinetic avidity breaks the trade-off between signal fidelity (requiring long half-life) and signal reversibility (requiring short half-life), which is a key requirement for T cell antigen discriminated mediated by kinetic proofreading.

## Introduction

Protein-protein binding domains are fundamental to signalling networks (1–3). Many binding domains recognise post-translational modifications - an archetypal example is the Src Homology 2 (SH2) domain, which binds to phosphorylated tyrosines within unstructured regions of proteins (4). SH2 domain-containing proteins are critical for signaling downstream of many surface receptors that become phosphorylated on their cytoplasmic tails upon ligand binding (e.g. receptor tyrosine kinases (5) and non-catalytic tyrosine-phosphorylated receptors (NTRs) (6)). Two well studied protein families that contain SH2 domains are Src and Syk kinases and in both families, SH2 domains are implicated in localisation and allosteric activation (7–9). Out of the 110 proteins in humans with SH2 domains, 100 contain a single SH2 domain (e.g. Src kinases) but only 10 contain tandem SH2 domains (e.g. Syk kinases). The precise function of tandem SH2 domains is unclear.

The Syk family contains two cytosolic proteins, Syk (Spleen tyrosine kinase) and ZAP70 (Zeta-chain-associated protein kinase 70), and both have tandem SH2 domains. They bind biphosphorylated immunotyrosine-based activation motifs (ITAMs, YxxL/Ix_6−8_YxxL/I) that are found on the cytoplasmic tails of activating NTRs, such as Fc receptors, B cell receptors, and T cell receptors (TCR) (6). Binding of tandem SH2 domains to biphosphorylated ITAMs is thought to improve specificity (10), increase affinity (11–13), and/or induce structural allosteric activation of the kinase domain (14–18). However, these functions are not unique to tandem SH2 domains raising the question of whether tandem domains simply have quantitative advantages over single SH2 domains or whether they can exhibit qualitatively distinct behaviours.

An often overlooked property of signalling networks is the mechanism(s) by which they can be efficiently reversed. In T cells, binding of the TCR to peptides presented on major histocompatibility complexes (pMHCs) can induce phosphorylation of ITAMs by the Src-family kinase Lck leading to the recruitment and subsequent phosphorylation of ZAP70 at the TCR (9). The delay between pMHC binding and the activation of ZAP70 is thought to contribute to the kinetic proofreading chain that is critical for the ability of T cells to discriminate between short-lived self and longer-lived non-self pMHC interactions (19–26). Critically, this mechanism requires that all signalling reactions within the chain are rapidly reversed upon pMHC unbinding so that short-lived self pMHC cannot exploit sustained TCR signals to short circuit the chain. Although highly active and abundant phosphatases, such as CD45, are known to dephosphorylate TCR signalling (27, 28), it is unclear how the high affinity ZAP70-ITAM interaction (11, 29, 30) can be reversed given that SH2 domains shield phosphotyrosines from phosphatases (31). This highlights a trade-off with high affinity SH2 domain interactions: they can maintain signalling of ligand bound receptors but, by preventing receptor dephosphorylation, they may allow receptors to sustain signalling even after ligand unbinding.

Here, we use modelling to show that kinetic proofreading can be maintained when TCR unbinding from pMHC selectively accelerates ZAP70 unbinding from the TCR. We show using modelling and experiments that this regulated unbinding can take place if ZAP70 binds to ITAMs by a kinetic, but not a static, avidity mode. A prediction of the model is that the ZAP70 half-life in T cells will be correlated to the TCR/pMHC half-life and we show that this is the case using live-cell microscopy. The work highlights that tandem SH2 domains can break the trade-off between signal robustness (requiring a long half-life) and signal reversibility (requiring a short half-life) to faithfully couple TCR/pMHC binding with TCR signalling, a key requirement for kinetic proofreading.

## Results

### Regulated ZAP70 unbinding can maintain kinetic proofreading at the TCR

A key requirement for kinetic proofreading is that the TCR rapidly resets to its basal state upon pMHC unbinding (Fig. 1A). This ensures that the TCR does not sustain any signal between pMHC binding events that would allow pMHC with faster off-rates to short-circuit the proofreading chain. Although phosphatases may rapidly shut off signalling by dephosphorylating exposed tyrosines, such as those on ZAP70 or ITAMs, it is less clear how ZAP70 recruitment can be rapidly reversed given that SH2 domains protect phosphorylated tyrosines (31). Modifying kinetic proofreading to explicitly include ZAP70 recruitment introduces a sustained signalling state whereby ZAP70 can remain bound to the TCR even after pMHC unbinding (Fig. 1B, *R*_2_). As a result of this sustained signalling, specificity produced by this model is reduced compared to a standard 3-step proofreading model, at all values of the ZAP70 unbinding rate *k*_off_ (Fig. 1C, *k*_reg_ = 0). In this model, ZAP70 remains bound to the TCR when *k*_off_ is small short-circuiting proofreading so that it effectively has only a single step (ZAP70 phosphorylation). On the other hand, ZAP70 is seldom bound when *k*_off_ is large so that the only robust proofreading step is ITAM phosphorylation and in this regime, T cell sensitivity to antigen is impaired (Fig. 1D). Hence, a trade-off emerges between signal fidelity and signal reversibility. Kinetic proofreading can be restored if ZAP70 unbinding can be selectively accelerated when the TCR is unbound from pMHC. Introducing this regulated unbinding (with rate *k*_reg_), we find that the proofreading chain can be restored with improved specificity when *k*_off_ *<*1 s^−1^ (Fig. 1C).

**Figure 1:**
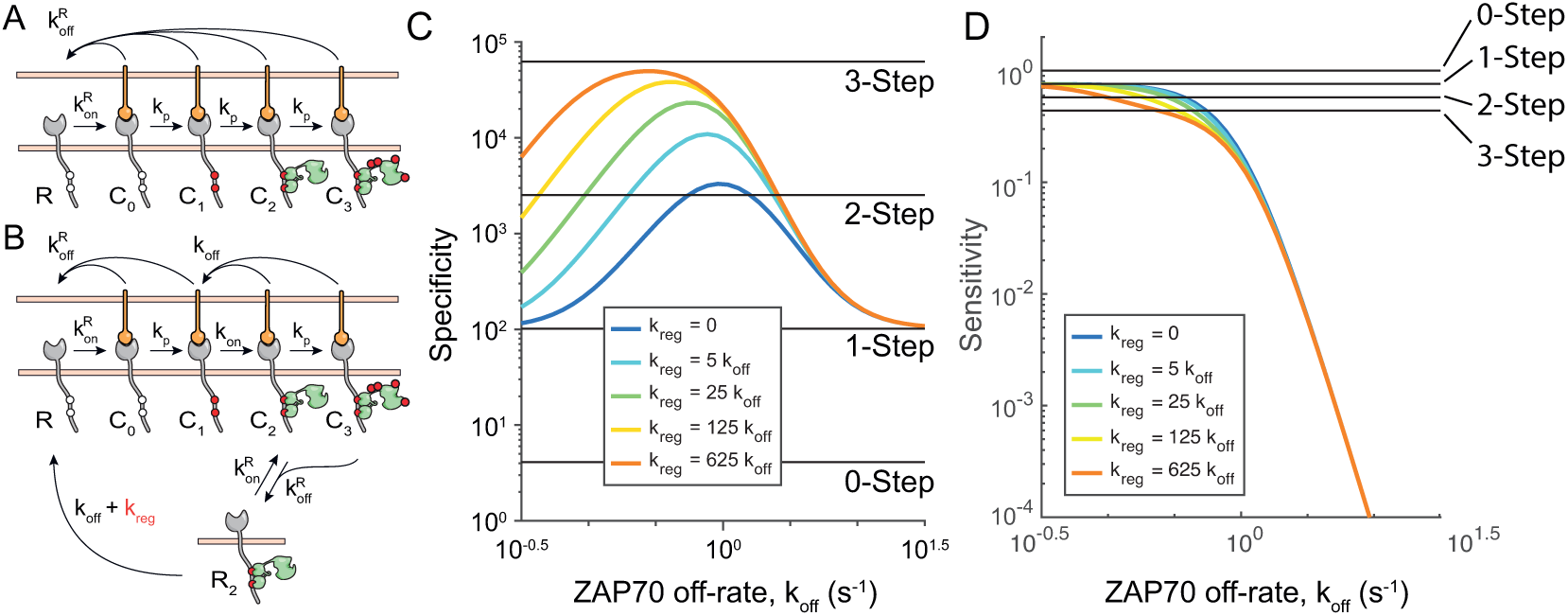
Operational model shows that ZAP70 unbinding abrogates kinetic proofreading unless it is regulated. (A) Standard 3-step kinetic proofreading showing that pMHC binding to the TCR initiates a sequence of steps (1 - ITAM phosphorylation, 2 - ZAP70 recruitment, 3 - ZAP70 phosphorylation) that can result in a signalling active TCR (*C*_3_). All steps are assumed to be immediately reversed upon pMHC unbinding 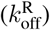. (B) Modified kinetic proofreading that explicitly models ZAP70 recruitment showing that ZAP70 can bind and unbind to phosphorylated TCR when it is bound to pMHC (with rate *k*_on_ and *k*_off_) and that ZAP70 can remain bound after pMHC unbinding (*R*_2_). Regulated unbinding (*k*_reg_) is introduced to selectively accelerate unbinding of ZAP70 when the TCR is unbound (state *R*_2_). As in the standard model, dephosphorylation is assumed to be immediate upon pMHC unbinding. (C) Specificity and (D) Sensitivity over the ZAP70 off-rate for the indicated values of the regulated off-rate *k*_reg_ (colours). Specificity is defined as the fold-change in *C*_3_ for a 100-fold change in the TCR/pMHC off-rate and sensitivity is a normalised value of *C*_3_ for the smallest value of the TCR/pMHC off-rate. Horizontal black lines are results for standard proofreading model with the indicated number of steps. See Methods for details.

### ZAP70 binding to ITAMs by kinetic, but not static, avidity supports regulated unbinding

The ZAP70/ITAM interaction may be approximated by a 1:1 binding model, which describes single SH2 domain interactions, if ZAP70 spends the majority of its time simultaneously engaged with both SH2 domains (Fig. 2A). We refer to this mode of binding as static avidity since binding is characterised by a single dominant state. We used this model to calculate the amount of bound ZAP70 over time with different phosphatase activities but with a constant concentration of solution ZAP70, as would be the case in T cells where ZAP70 in the cytoplasm can continuously bind and unbind to phosphorylated ITAMs (Fig. 2B). The observed unbinding rate (i.e. the rate at which ZAP70 occupation of ITAMs decays to zero) increased with increasing phosphatase activity but was limited to the ZAP70 off-rate since phosphatases cannot dephosphorylate ITAMs bound by SH2 domains (Fig. 2E).

**Figure 2:**
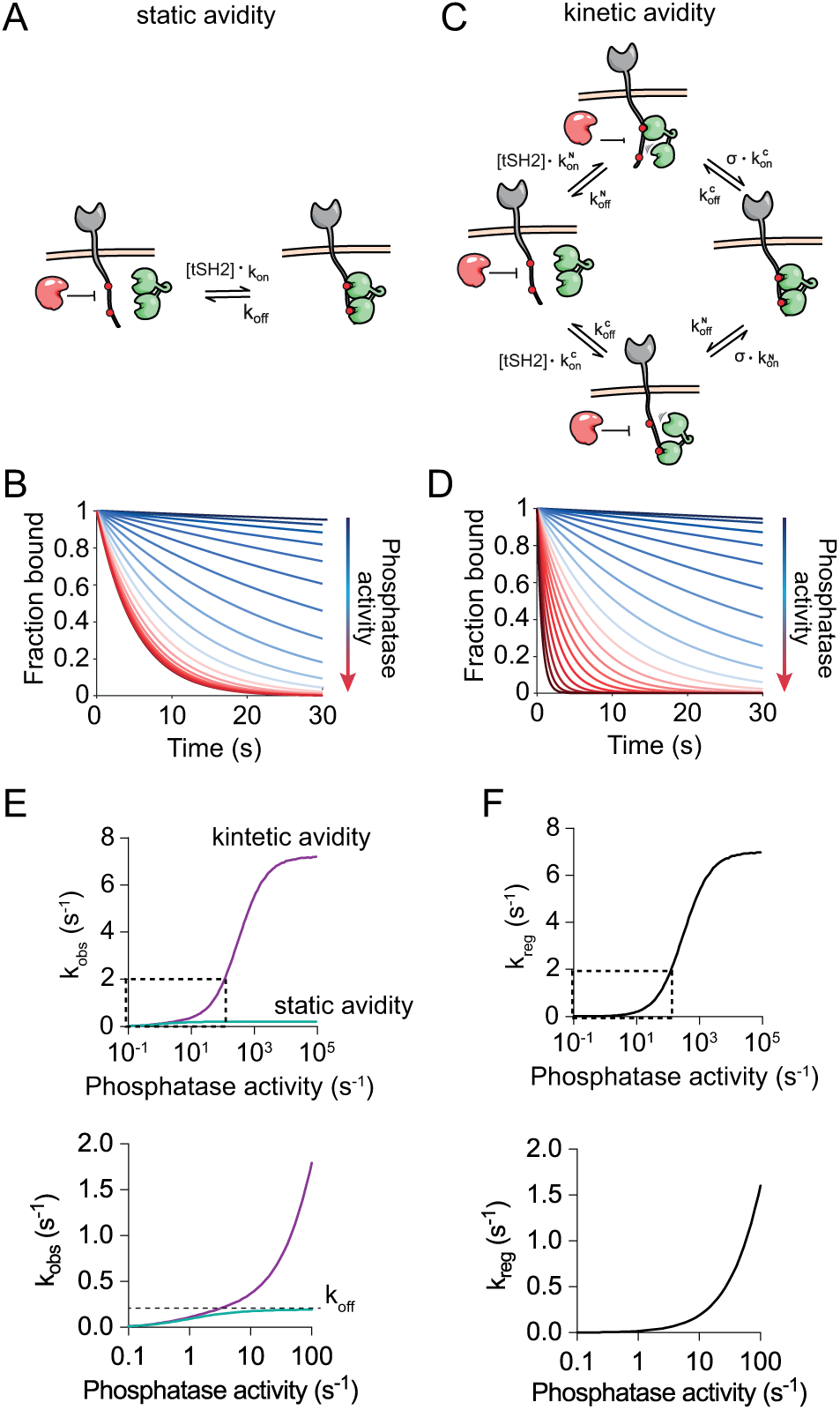
ZAP70 binding to ITAMs by kinetic but not static avidity predicts regulated unbinding by phosphatases. (A) A standard 1:1 binding model showing that phosphatases (red) can dephosphorylate exposed (non-bound) phosphotyrosines. (B) Increasing phosphatase activity increases the observed unbinding rate of SH2 domains. (D) Kinetic avidity model for ZAP70 parameterised by an on-rate and off-rate for each SH2 domain binding each phosphotyrosine and by the local concentration of phosphotyrosine experienced by the unbound SH2 domain when ZAP70 is bound by the other SH2 domain (*σ*). In this model, phosphotyrosines are exposed even when ZAP70 is bound to the ITAM. (D) Increasing phosphatase activity increases the observed unbinding rate beyond the ZAP70 *k*_off_. (E) The observed unbinding rate over PTP activity determined by exponential fit to static (green) and kinetic (purple) avidity models. (F) The regulated unbinding rate calculated by the difference in observed off-rates between the kinetic and static avidity calculations in panel E.

We next constructed a detailed ZAP70 binding model (Fig. 2C), where we explicitly included the initial interaction of each SH2 domain and the subsequent interaction of the second domain mediated by intrinsic on-rates that are dependent on the effective concentration of free phosphotyrosine (which we refer to as *σ*, in units of *µ*M). Using model parameters that allowed ZAP70 to rapidly cycle between these internal states, yet achieve the same overall unbinding rate used in the simpler 1:1 binding model above, we found that the observed off-rate could exceed the ZAP70 off-rate (Fig. 2D,E). By taking the difference in observed off-rates between the models, we directly calculated the contribution of regulated unbinding resulting from phosphatases dephosphorylating ITAMs interacting with ZAP70 and parsing it from dephosphorylation of free ITAMs (Fig. 2F). Therefore, ZAP70 binding to ITAMs by kinetic avidity supports regulated unbinding by phosphatases.

### Systematic analysis predicts kinetic avidity between ZAP70 and ITAMs

To determine the binding mode of ZAP70, we experimentally determined all model parameters. To do this, we used the tandem SH2 domains of ZAP70 (tSH2) and bi- or monophosphorylated ITAM peptides derived from the membrane-distal ITAM of CD3*ζ* (ITAM3). As expected, the 1:1 binding model was able to describe binding of tSH2 to monophosphorylated ITAM3 N and C peptides at steady-state (Fig. 3A, B). A kinetic analysis showed that on-rates were not measurable by SPR due to mass transport, however estimates of the off-rates were possible revealing rapid unbinding (Fig. 3C, D). Therefore, on-rates were determined using off-rates and dissociation constants (Table 1).

**Table 1:** Affinity and kinetic measurements of ZAP70 on ITAM3 peptides by surface plasmon resonance (SPR) at 37°C (N ≥3). − values estimated from measured *K*_D_ and *k*_off_.

| Peptide | $K_D$ ($\mu\text{M}$) | $k_{\text{on}}$ ($\mu\text{M}^{-1} \text{s}^{-1}$) | $k_{\text{off}}$ ($\text{s}^{-1}$) | $t_{1/2}$ (s) |
| --- | --- | --- | --- | --- |
| ITAM3 N | $20.7 \pm 2.7$ | 0.265* | $5.5 \pm 0.46$ | 0.126 |
| ITAM3 C | $7.52 \pm 1.0$ | 1.52* | $11.4 \pm 4.6$ | 0.061 |
| ITAM3 NC | $0.0962 \pm 0.024$ | $1.89 \pm 0.356$ | $0.227 \pm 0.036$ | 3.05 |

**Figure 3:**
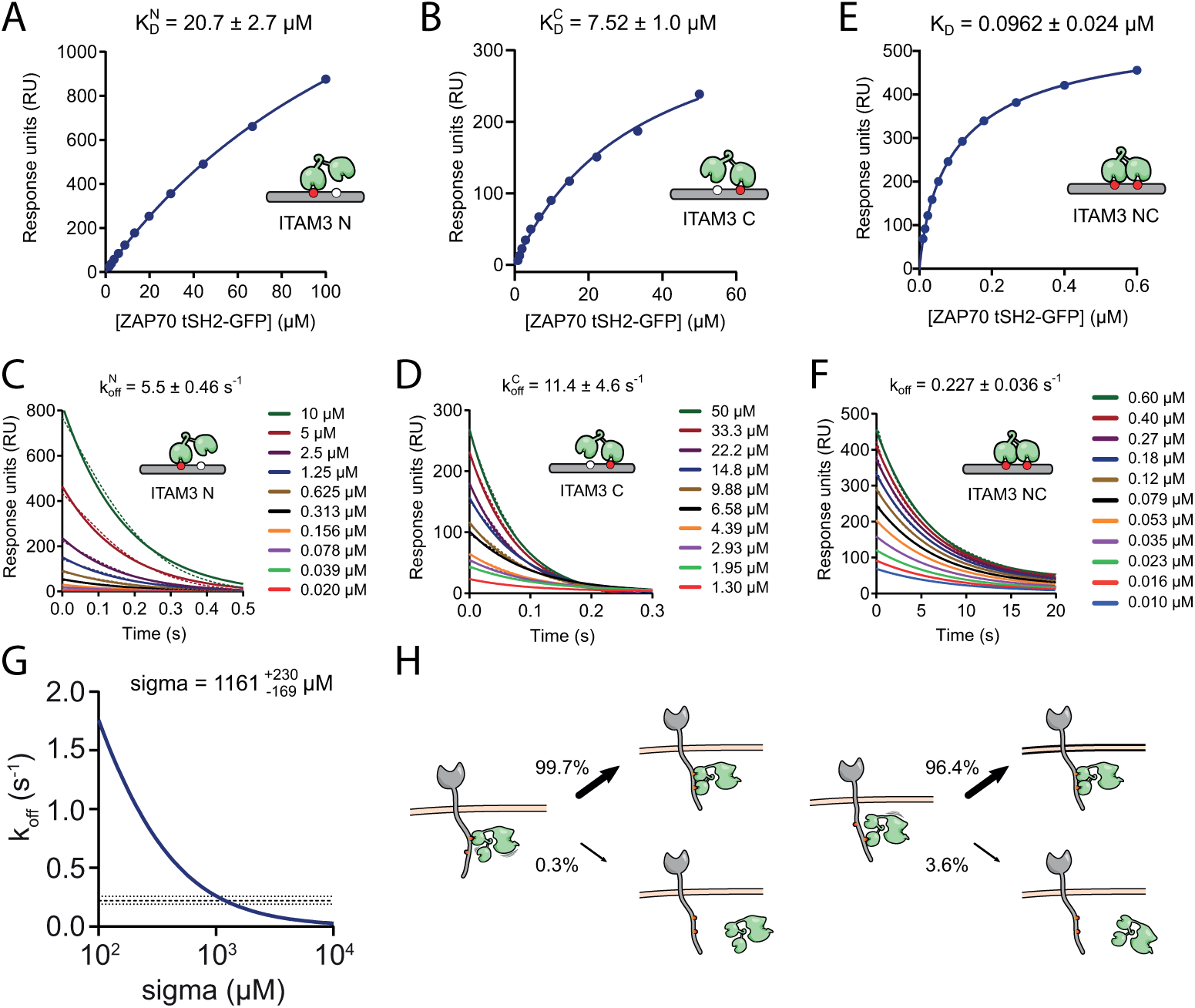
SPR binding affinity and kinetics of ZAP70 interacting with mono- and biphosphorylated ITAM peptides parameterise the mathematical model and predict kinetic avidity. Representative equilibrium and dissociation phase data from tSH2-GFP protein interacting with (A,C) N- or (B,D) C-monophosphorylated or (E,F) biphosphorylated CD3*ζ* ITAM3 peptides. Indicated parameters are obtained by fitting the data (dots) with a 1:1 binding model (solid lines). (G) Unbinding rates of ZAP70 from biphosphorylated ITAM (y-axis) were calculated using the model for different values of *σ* (x-axis) with the 4 kinetic rate constants fixed to their experimentally determined values (Table 1). The mean (dashed line) ±SEM (dotted lines) of the bivalent dissociation rate are shown. (H) Probability of rebinding or unbinding when ZAP70 is bound to N-terminal (left panel) or C-terminal (right panel) tyrosines of ITAM3 using the value of *σ* determined in (G) (see main text for calculation) predicts that the long half-life of ZAP70 is achieved by kinetically cycling between states bound by one and both SH2 domains.

The interaction between tSH2 and biphosphorylated ITAM peptides were also, unexpectedly, well described by a 1:1 binding model revealing an effective *K*_D_ of 0.096 *µ*M (Fig. 3E), *k*_off_ of 0.227 s^−1^ (Fig. 3F), and *k*_on_ of 1.89 *µ*M^−1^s^−1^ (Fig. S1). Although unexpected, this was predicted by the bivalent model, which produced a binding equation that was identical in form to the monomeric model except that the fitted *K*_D_ value was dependent on the individual reaction parameters: 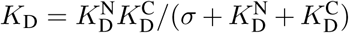 (see Methods).

Given that all dissociation constants were determined, it was possible to directly calculate *σ* to be 1733 ± 743 *µ*M. We also estimated *σ* directly from the kinetic data by calculating the ZAP70 off-rate for different values of *σ* fixing the 4 kinetic rate constants to their experimentally determined values. As expected, the predicted ZAP70 off-rate decreased when *σ* increased with the experimentally determined off-rate obtained when *σ* was 1161 *µ*M (Fig. 3G), which was within the error of the value determined above and similar to a recent theoretical estimate for Syk (32). Given that *k*_off_ is estimated with higher accuracy compared to *K*_D_ in SPR, we proceeded to use the estimate of *σ* obtained by the kinetic data.

Using this estimate of *σ*, we calculated the fraction of times that ZAP70, when bound by a single SH2 domain, would rebind with the other SH2 domain versus unbinding back to solution (e.g. 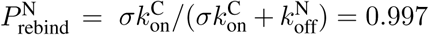, when ZAP70 is bound to ITAM N and 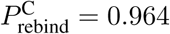, when ZAP70 is bound to ITAM C). These calculations revealed that ZAP70 rebinds over 96% of the time compared to unbinding from the ITAM showing that the long half-life is maintained by kinetic avidity whereby ZAP70 cycles between internal binding states (Fig. 3H).

A long half-life could also be achieved by static avidity, whereby the values of 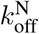 and 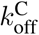 are specifically reduced when ZAP70 is simultaneously bound to both ITAM tyrosines so that binding is dominated by this single state. To examine this possibility, we modified the model to include a conformational cooperativity parameter (*λ*) that multiplied the off-rates only when ZAP70 is bound to both ITAM tyrosines (Fig. S2A). Using a similar fitting strategy we were unable to uniquely determined *λ* and *σ*. However, we were able to place bounds on them showing that the largest degree of conformational cooperativity (i.e. smallest value of *λ*) that could explain the data was 0.3 and at this value, *σ* was 300 *µ*M (Fig. S2B). With these parameters, we found that 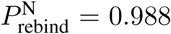 and 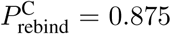 (Fig. S2C). This result highlights that even in a model with conformational cooperativity, rebinding is substantial indicating that kinetic avidity is responsible for the long half-life.

### The phosphatase CD45 can regulate the unbinding of ZAP70 from ITAMs

A key consequence of kinetic avidity is the ability of phosphatases to regulate unbinding, which we directly tested using an SPR assay for phosphatase-accelerated unbinding. In this assay an N-terminally Avitagged, biotinylated and phosphorylated peptide corresponding to the full intracellular tail of CD3*ζ* with all tyrosines mutated to Ala except the C-terminal ITAM (Avi-CD3*ζ* ITAM3) was immobilised on the chip surface. A near-saturating concentration of ZAP70 tSH2 was injected over the surface and allowed to reach steady-state before the same concentration of ZAP70 mixed with different concentrations of the cytoplasmic domain of the phosphatase CD45 were injected (Fig. 4A). This assay format allowed us to assess the rate of ITAM dephosphorylation by CD45 in the presence of a constant concentration of ZAP70, as would be the case in T cells.

**Figure 4:**
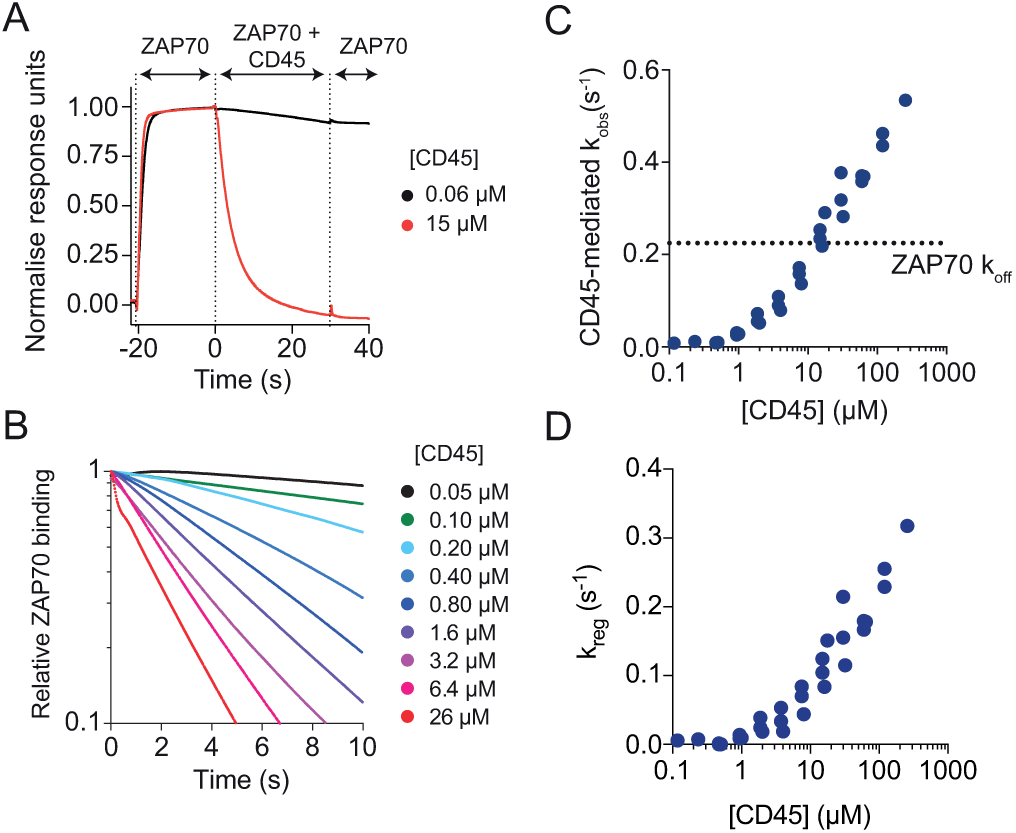
CD45 increases the unbinding rate of ZAP70 from ITAMs beyond the ZAP70 off-rate. (A) Example sensogram of CD45-accelerated ZAP70 unbinding assay in SPR: ZAP70 (500 nM) was first injected over a surface of ITAM3-phosphorylated CD3*ζ* cytoplasmic domain and allowed to reach steady-state, before a mixture of ZAP70 (500 nM) and CD45 (indicated concentration) was injected, and finally ZAP70 (500 nM) was injected. (B) ZAP70 and CD45 coinjection phase for multiple concentrations of CD45 demonstrating a concentration dependent acceleration in the loss of ZAP70 binding. (C) The fitted observed unbinding rate over [CD45] (results from three experiments conducted on different days are shown). The ZAP70 off-rate is shown as a dotted line. (D) The regulated off-rate calculated over [CD45]. Binding of ZAP70 to this full length Avi-CD3*ζ* ITAM3 was the same as on the shorter ITAM3 peptide (see Fig. S4).

In the presence of CD45, the amount of ZAP70 binding to the chip surface decreased over time (Fig. 4A,B). If CD45 only dephosphorylated free ITAMs, then the maximum observed rate of ZAP70 unbinding would be *k*_off_. However, the results clearly demonstrate that beyond 10 *µ*M of CD45 the observed off-rate exceeded *k*_off_ indicating that CD45 can dephosphorylate ITAMs bound by ZAP70 (Fig. 4B, C). As before, we calculated the CD45-mediated regulated off-rate by subtracting the experimental observed off-rate from the predicted observed off-rate in the static avidity model (Fig. 4D). This result was also observed with different immobilisation levels of Avi-CD3*ζ* ITAM3 demonstrating that SPR transport artefacts were unlikely to be affecting the results (Fig. S3A-D). Given that all ZAP70 binding parameters were identified, we were able to use this data to estimate that CD45 has a high catalytic efficiency of 0.103±0.01 *µ*M^−1^ s^−1^ for ITAM3 (Fig. S3E).

Although ZAP70 bound with the same affinity and kinetics to the ITAM3 peptide used in the previous section and the full length Avi-CD3*ζ* ITAM3 used here (Fig. S4), there seemed to be steric limitations to regulated unbinding from the shorter ITAM3 peptide since the CD45-mediated observed off-rate hit a plateau at the ZAP70 off-rate (Fig. S5). Since there are only 5 amino acids between the biotin and first phosphotyrosine in the ITAM3 peptide, a likely explanation is that there is not enough space to accomodate both ZAP70 and CD45 near the streptavidin anchor. Consistent with this, we have previously observed strong steric hindrance of phosphatase domains accessing phosphotyrosines immobilised on SPR chip surfaces with short linkers (33).

### The membrane half-life of ZAP70 in T cells correlates with the TCR-pMHC half-life

Having established that CD45 can regulate the unbinding of ZAP70 from ITAMs we sought evidence for this in T cells. We hypothesised that if pMHC unbinding exposes the TCR to phosphatases (28), then the half-life of ZAP70 at the membrane should correlate with the half-life of TCR-pMHC interactions. On the other hand, if the half-life cannot be regulated then only the number of ZAP70 molecules at the membrane would correlate with the TCR-pMHC half-life.

To test this hypothesis, we first established conditions for accurate off-rate measurements using imaging. We immobilised ITAM3 peptides on glass coverslips and used total internal reflection fluorescence (TIRF) microscopy together with single particle tracking (SPT) to measure the binding time of ZAP70 tSH2-GFP or ZAP70 tSH2-Halotag-Alexa647. By fitting the distribution of ZAP70 binding times to exponentials, we were able to determine the off-rate for monophosphorylated ITAM3 N (11.4±0.011 s^−1^) and ITAM C (4.75±0.14 s^−1^), and biphosphorylated ITAM NC (0.368±0.011 s^−1^) peptides (Fig. S6, see Methods). These kinetic rate constants were in excellent agreement with those determined by SPR validating the imaging conditions.

We next used supported lipid bilayers with biotinylated pMHC and ICAM-1 to stimulate T cells and monitored the recruitment of fluorescent ZAP70 constructs (Fig. 5A). Full length ZAP70-Halotag was introduced into ILA TCR expressing Jurkat cells with normal endogenous expression of ZAP70 and labelled with low levels of Alexa647-Haloligand so that single molecules of ZAP70 recruited to the membrane could be monitored with SPT (Fig. 5B). As expected, the amount of membrane-recruited ZAP70 showed a strong correlation with the TCR-pMHC half-life (Fig. 5C and D). Notably when pMHC was substituted for anti-CD90 in the bilayer to promote adhesion without TCR engagement, we still observed some residual recruitment of ZAP70 (Fig. 5C), consistent with the observation of ligand-independent TCR triggering (38–40).

**Figure 5:**
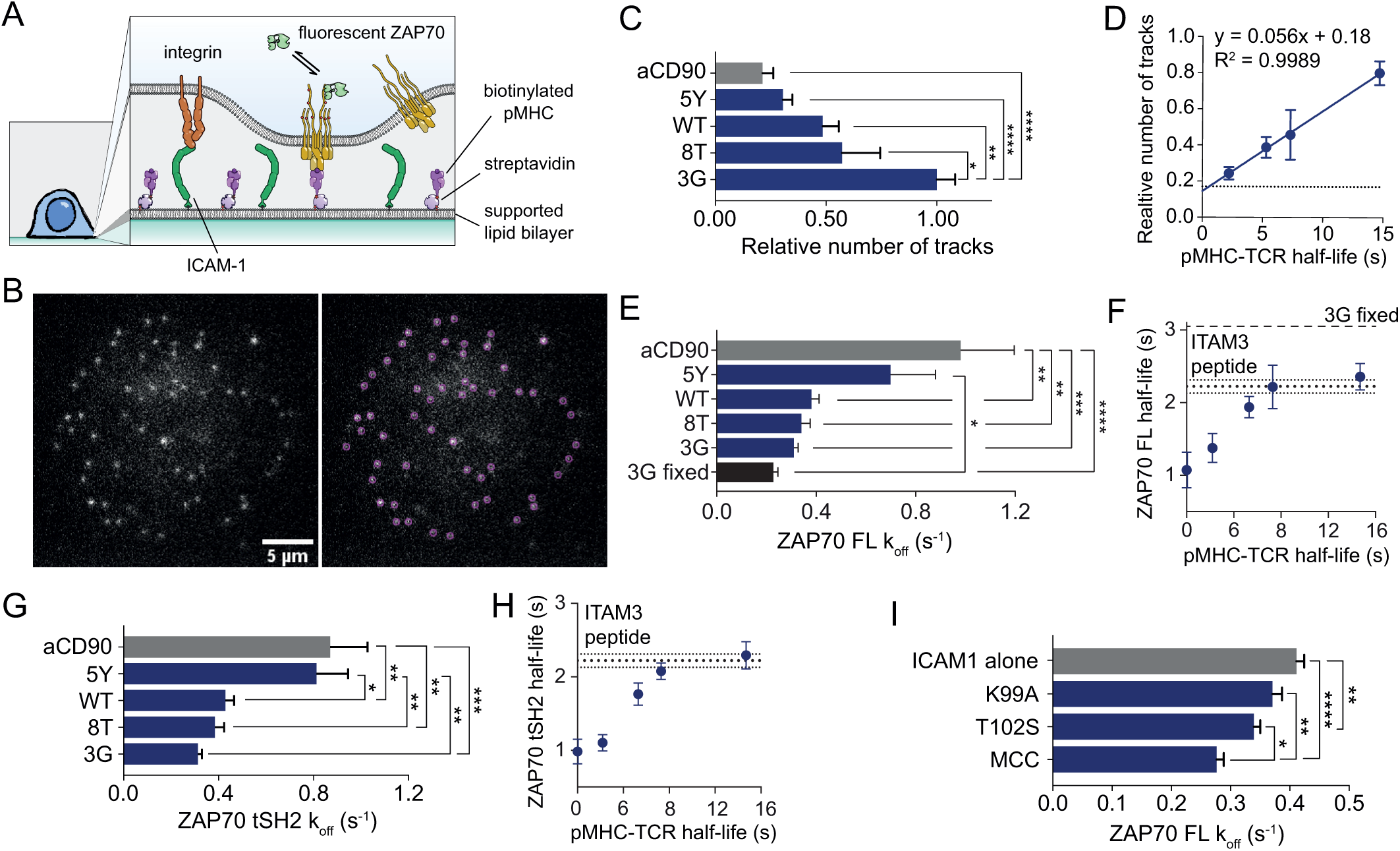
ZAP70 membrane half-life correlates with the TCR-pMHC half-life in T cells. (A) Diagram of experimental system. (B) Example frame (left panel) and identified particles (right panel) of a live-cell ZAP70-Halotag SPT experiment with 3G pMHC and ICAM-1. (C) Number of labelled ZAP70 recruited to the interface between ILA Jurkats and pMHC-decorated SLBs normalised to the highest affinity pMHC (3G). (D) Number of ZAP70-Halotag over the TCR-pMHC half-life (measured at 25°C (37)), with the horizontal line indicating the anti-CD90 condition. (E) Fitted *k*_off_ and (F) half-life calculated from *k*_off_ over the TCR-pMHC half-life from the distribution of membrane binding times. (G,H) Repeat of experiments in (E,F) except with the truncated tSH2-GFP instead of full length ZAP70-Halotag. (I) The *k*_off_ of full length ZAP70-GFP recruited to the membrane at the interface of primary AND TCR transgenic mouse CD4^+^ T cells in live-cell SPT experiments. All binding time distributions were fit with a sum of two exponentials with the slow rate displayed (see Methods for details). Data are from at least 8 cells per condition imaged in three separate experiments. Mean ±SEM are shown, * p*<*0.05, ** p*<*0.01, *** p*<*0.001, **** p*<*0.0001 (one-way ANOVA with Tukey’s post-test).

In support of our hypothesis, measurements of the ZAP70 membrane half-life also correlated with the TCR-pMHC half-life (Fig. 5E and F). As the TCR-pMHC half-live increased, we observed that the membrane half-life of ZAP70 appeared to reach a plateau at the rate we measured with isolated peptide and recombinant proteins (0.310 s^−1^ for 3G in Fig. 5E, F compared to 0.368 s^−1^ in Fig. S 6D). The plateau was not a technical limit because trapping ZAP70 at the membrane by stimulating T cells with 3G and fixing prior to imaging showed a slower off-rate (3G fixed, Fig. 5E,F). Although we detected recruitment of ZAP70 without TCR engagement, it dissociated more rapidly (Fig. 5E, anti-CD90).

Recently, it has been suggested that Lck-mediated phosphorylation of interdomain B tyrosines in ZAP70 increase the half-life at the TCR (18). To control for this non-exclusive mechanism, which would also theoretically be regulated by phosphatases, we used the same tSH2-GFP construct used in SPR experiments. Despite lacking interdomain B, tSH2-GFP completely reproduced the results from full length ZAP70-Halotag protein (Fig. 5G, H), suggesting that kinetic avidity is sufficient to explain how the ZAP70 half-life is responsive to the TCR-pMHC half-life.

Finally, we reproduced the results in primary AND TCR transgenic mouse CD4^+^ T cells presented with a set of altered peptide ligands. Although the kinetics are not known, functional data demonstrates that the index MCC peptide more robustly activates T cells compared to the T102S and K99A altered peptide ligands (41). In this system we again observed recruitment of ZAP70 without engagement of TCR, this time with only ICAM-1 in the bilayer (Fig. 5I). In agreement with the Jurkat results, we found significant differences in the ZAP70 membrane off-rate (Fig. 5I), supporting the hypothesis that the regulated ZAP70-ITAM half-life is sensitive to TCR-pMHC half-life.

It has been reported that the half-life of SH2 domains at the membrane can be up to 20-fold longer than their *in vitro* measured half-life by a rebinding mechanism (42). By applying the analysis used by Oh et al to our data we found no evidence for ZAP70 rebinding (Fig. S8); the SPR and membrane half-lives were similar (Fig. S8A) and the membrane diffusion coefficient of ZAP70, for pMHCs with different ZAP70 half-lives, did not correlate with its half-life (Fig. S8B,C). The rebinding mechanism was shown to require a high density of free binding sites (∼1000 sites / *µ*m^2^) distributed on the *µ*m scale where SH2 domains rebound across different receptors (42). This is unlikely to be the case for T cells where low pMHC densities generate spatially restricted TCR clusters. Moreover, stochastic spatial simulations show that rebinding between the same ZAP70 and the same TCR is unlikely even if the TCR maintains 100 free ITAMs (Fig. S7). This result is consistent with previous studies showing that rebinding to the same spatially-localised site is not significant for cytosolic diffusion coefficients of *D* ∼ 10 *µ*m^2^/s (although appreciable for typical membrane diffusion coefficients of *D* ∼ 0.05 *µ*m^2^/s) (43–45).

## Discussion

The tandem SH2 domains of ZAP70 were previously shown to increase affinity and specificity, and to contribute to the allosteric activation of the kinase (9). Here, we report an additional feature; the ability to exhibit regulated unbinding to ITAMs.

The mechanism of kinetic avidity for ZAP70 is likely to be influenced by flexibility in the ITAM sequence. In ZAP70, the tandem SH2 domains are connected by a short coiled-coil region and the N-terminal SH2 binding site includes some residues from the interface with the C-terminal SH2 domain (14, 46). These structural observations have suggested that the SH2 domains are locked together with little flexibility upon ITAM binding (9), but the unstructured ITAM is likely to remain flexible so that individual phosphotyrosines can move away from either SH2 domain upon unbinding. Thus, a highly kinetic binding mode can take place even if ZAP70 is relatively rigid.

It is noteworthy that previous *in vitro* measurements of the ZAP70 half-life are significantly longer than those reported here. In early SPR measurements (11, 29), the tSH2 of ZAP70 did not unbind on the timescale of 100 s, suggesting *k*_off_ *<* 0.01 s^−1^ and this may be a consequence of the lower temperatures used in those experiments where affinities are known to be higher. A more recent report (30) using Bio-Layer Interferometry (BLI) reported *k*_off_ ∼ 10^−4^ s^−1^ (half-life of 1.9 hours) for this interaction. Given that BLI is an optical technique that requires large amounts of protein binding for detection, the high amount of peptide immobilisation used to achieve this may mean that a large amount of intermolecular rebinding across different ITAMs is taking place leading to apparent long half-lives. Consistent with the present work, previous *in vivo* measurements of the ZAP70 half-life using fluorescence recovery after photobleaching have provided recovery timescales of ∼10 s (18, 47, 48).

How does regulated unbinding relate to established mechanisms of ZAP70 activation? Detailed analyses have shown that ZAP70 recruitment is accompanied by alignment of the SH2 domains that release ZAP70 from autoinhibition allowing for phosphorylation that increases its activity (16, 18, 49, 50). As discussed above, it is likely that the cycling of ZAP70 between states that expose ITAM tyrosines takes place while the SH2 domains remain aligned allowing for ZAP70 phosphorylation. Under this assumption, we formulated a detailed molecular model of ZAP70 activation that explicitly includes ITAM and ZAP70 phosphorylation when the TCR is bound and dephosphorylation otherwise. These reactions were coupled to the fully-parametrised model of ZAP70 binding dependent on ITAM phosphorylation but not on TCR/pMHC binding (Fig. S9), and in this way, regulated unbinding was a consequence not an assumption of the model. In this model, higher specificity was observed with higher phosphatase activity and with ZAP70 binding by kinetic rather than static avidity. The binding of ZAP70 to the TCR was observed under all conditions but only pMHC binding to the TCR with lower off-rates were also able to activate ZAP70, which is consistent with previous reports (51, 52). This observation is a result of ZAP70 binding being an earlier step in proof-reading so that recruitment exhibits less specificity compared to downstream activation and this has recently been observed using a chimeric antigen receptor system (53). Finally, the ‘catch and release’ mechanism of ZAP70 activation (54) is consistent with the present work but we note that if released and activated ZAP70 is caught by different TCRs, it will short-circuit proofreading reducing antigen discrimination. However, this is unlikely given that ZAP70 rebinding to different TCRs does not seem to be taking place (Fig. 8).

Finally, we note that regulated unbinding may be pervasive. There are 9 other proteins with tandem SH2 domains and a large number of proteins with domains that bind regulated sites (i.e. a site that can either be on or off), which include SH2, PTB, FHA, and PH domains. The qualitative feature of regulated unbinding cannot take place with 1 domain and it does not require more than 2. Interestingly, out of 447 proteins that contain these domains, there are no proteins that contain more than 2 domains per protein with the exception of 3 PH domain containing proteins (Fig. S10). In contrast, constitutively binding domains, such as SH3, PDZ, C2, and WW domains, are found in copy numbers that exceed 2 on 66 proteins (Fig. S10). In addition to acting between two proteins, regulated unbinding is likely to operate on stable phase-separated signalling assemblies formed by weakly binding multi-domain proteins (55, 56). Interestingly, activated ZAP70 catalyses the formation of LAT signalling assemblies (57) and recently, it has been suggested that LAT may be a proofreading step (58, 59). Consistent with this, *in vitro* generated LAT assemblies displayed long half-lives without phosphatases but were disassembled within minutes in their presence (57). Future work is needed to examine the regulated disassembly kinetics of LAT within intact T cells.

## Acknowledgements

We thank Oreste Acuto, Johannes Huppa, Daniel Coombs, and Marion H. Brown for helpful discussion. This work was supported by the Wellcome Trust (SIA 101799/Z/13/Z to AvdM, PRF 100262Z/12/Z to MLD, SRF 207537/Z/17/Z to OD), European Commission (ERC-2014-AdG 670930 to MLD), the Kennedy Trust for Rheumatology Research (MLD), the Medical Research Council (MR/J002011/1 to OD and AvdM), the National Health and Medical Research Council of Australia (APP1163814 to JG), and the National Science Foundation, USA (NSF-DMS 1902854 to SAI).

## Methods

### Plasmids and peptides

For bacterial expression, a construct of the tandem SH2 domains of human ZAP70 (amino acids 1-264) and a C-terminal GFP or Halotag was cloned into pET21 between the NdeI and NotI restriction sites. For SPR experiments the cytoplasmic domain of CD3*ζ* (amino acids 52-164) with an N-terminal Avitag and a C-terminal hexahistadine tag was constructed by cloning the an Avitag-CD3*ζ* fusion into pET21 between the NdeI and NotI sites. For viral transduction full length human ZAP70 fused to Halotag were cloned into pHR-SIN-BX between the BamHI and NotI sites, removing the IRES-Emerald sequence. For *in vitro* transcription full length mouse ZAP70-GFP fusion was cloned into pGEM64 between AgeI and NotI.

All peptides were ordered from PeptideSynthetics and were certified to be *>*95% pure. Sequences of peptides used were ITAM3 N (biotin-GKGHDLLY*QGLSTATKDTYDALHMQ), ITAM3 C (biotin-GKGHDLLYQGLSTATKDTY*DALHMQ) and ITAM3 NC (biotin-GKGHDLLY*QGLSTATKDTY*DALHMQ), where Y* denotes a phopshotyrosine.

### Protein production

ZAP70 tSH2-GFP, ZAP70 tSH2-Halotag, CD45 catalytic domain DNA constructs were transformed into the BL21 (DE3) strain (NEB) Escherichia coli and plated on LB agar with ampicillin (100 mg/ml), and then grown overnight at 37°C. For the avitag-CD3*ζ* ITAM3 peptide DNA construct BirA-transformed BL21 (DE3) Escherichia coli (BPS Bioscience) were used. The next day, colonies were innoculated into a 10 ml LB selection medium (LB medium with 100 mg/ml ampicillin), grown overnight at 37°C, and then transferred to 1 liter of LB selection medium until the optical density at 600 nm was 0.6 to 0.8. The cells were then treated with isopropyl-1-thio-D-galactopyranoside (final concentration, 0.5 mM) and harvested by centrifugation after 20 hours of culture at 18°C.

Bacterial pellets were resuspended in tris-buffered saline (TBS; 20 mM tris(hydroxymethyl)aminomethane, 150 mM NaCl) with protease inhibitors (protease inhibitor cocktail; Sigma), and then lysed with three 30 s bursts of sonication interspersed with 60 s rest periods on ice. Lysates were clarified by centriguation at 15,000 rcf followed by filtration through a 0.22 *µ*m filter. Clarified lysates were applied to Ni^2+^-NTA resin, which was washed with 10 column volumes of 2x PBS (10 mM phosphate, 300 mM NaCl), followed by 10 column volumes of 2x PBS with 10 mM imidazole, before His-tagged protein was eluted with 10 ml of 100 mM imidazole (pH 7.5). Protein was concentrated to 1 ml using a Amicon Ultra-4 Ultracel 30 kDa spin concentrator (Merck Millipore) and the protein was separated on a HiPrep 16/60 Sephacryl S-200 HR column (GE Healthcare) equilibrated with TBS. Peaks corresponding to the protein of interest were pooled, glycerol was added to a final concentration of 10% (v/v), and protein was stored in aliquots at 80°C until the day of experiment.

Avitag-CD3*ζ* ITAM3 was further phosphorylated *in vitro* using purified Lck in kinase buffer (TBS with 2 mM Mg_2_SO_4_, 500 *µ*M ATP and 1 mM DTT) for 1 hour at room temperature.

On the day of the experiment, aliquots of proteins were thawed and buffer exchanged into 20 mM HEPES, 150 mM NaCl, 1 mM EDTA, 0.005% Tween 20, and 1 mM dithiothreitol using a Zeba 7kDa spin column (Thermo Fischer). Concentrations of proteins were measured using the optical density at 280 nm, or 490 nm for GFP tagged proteins, using a DeNovix DS-11 FX spectrophotometer (Denovix).

### Surface Plasmon Resonance

Experiments were performed on a Biacore S200 instrument (GE Healthcare Life Sciences). All experiments were performed at 37°C using HEPES-buffered saline (HBS-EP, 10 mM HEPES (pH 7.4), 150 mM NaCl, 1 mM EDTA, and 0.005% Tween P20) and a flow rate of 50 *µ*l/min.

CM5 sensor chips were coupled with streptavidin to near saturation (typically between 4000 and 7000 RU) using the amine coupling kit (GE Healthcare Life Sciences). After streptavidin was coupled, biotinylated peptides were injected to give the indicated concentrations in experimental flow cells. For kinetic and equilibrium binding experiments excess biotin-binding sites were blocked with biotin in HBS-EP. Reference flow cells were treated with buffer and then blocked with biotin. The chip surface was then conditioned with 5 injections of HBS-EP followed by the indicated concentrations of ZAP70 constructs. Reference-subtracted data are shown.

For CD45 assays flow cells were not blocked with biotin after Avitag-CD3*ζ* ITAM3 peptide immobilisation. To perform the phosphatase assay an ABA injection method was used with ZAP70 tSH2-GFP (500 nM) as the flanking solution for an injection of a mixture of ZAP70 (500 nM) and CD45 at various concentrations. The flanking solution was first injected over a surface of Avitag-CD3*ζ* ITAM3 for 20 seconds to allow ZAP70 binding to reach equilibrium, then the mixture of ZAP70 tSH2-GFP and CD45 was injected. After each cycle a 30 s injection of 1x PhosSTOP (Merck) phosphatase inhibitors was performed and fresh biotinylated Avitag-CD3*ζ* ITAM3 immobilised.

### Cells

Jurkat T cells (Clone E6.1, ATCC) were maintained in RPMI 1640 (Gibco) supplemented with 10% (vol/vol) FCS, 2 mM L-glutamine, and 1 mM penicillin and streptomycin (all from Invitrogen). Jurkat cells stably expressing the ILA TCR were used in this study. The ILA TCR recognizes residues 540-548 (ILAKFLHWL) of the human telomerase reverse-transcriptase protein, presented in the context of the human MHC class I HLA-A*0201 (HLA-A2). To activate the ILA expressing Jurkat cells, monomeric, biotinylated human HLA-A2 complexed with the ILA peptide and several altered peptides (3G, 8T, 5Y) were used.

ZAP70-tSH2 GFP protein was introduced into ILA Jurkat cells by directly electroporating the purified protein (NEON; Invitrogen). Full-length ZAP70-Halotag was transduced into Jurkat cells using a lentiviral system and were labelled with Halotag ligand-Alexa647 made in house by reacting Halotag Amine ligand (Promega) with excess Alexa647-succinimidyl ester (Thermo Fischer) and neutralising excess reactive dye with glycine. To label cells 1 *µ*M of Halotag ligand-Alexa647 in complete culture media was incubated with cells for 30 min at 37°C, then the cells were washed twice with 20 ml of PBS, incubated at 37°C in complete media for 30 min, washed once more in PBS before imaging.

Primary CD4+ T cells were isolated from spleens of AND TCR transgenic mice using magnetic bead sorting. Messenger RNA encoding mouse ZAP70-GFP was transcribed *in vitro* and introduced into cells by electroporation 16 hours before imaging.

All cells were imaged in phenol-free RPMI with 10% FCS and 10 mM HEPES pH 7.5.

### Total internal reflection microscopy and particle tracking

Supported lipid bilayers were created and loaded with pMHC and His-tagged ICAM-1 as described elsewhere (61). Live cell image sequences were acquired on a total internal reflection fluorescence (TIRF) microscope (ELYRA, Zeiss) with an environmental chamber equilibrated to 37°C. Chamber-slides with pMHC-decorated bilayers were equilibrated in the microscope environmental chamber at least 10 minutes before cells were added to wells and imaging commenced as soon as cells landed on the coverslip. Cells were imaged within 10 min of addition to the bilayer-containing wells.

GFP imaging was performed with 0.7 mW of 488 nm light and Alexa647 was imaged with 0.5 mW of 647 nm light. A 100 oil-immersion objective (NA = 1.46) was used for all imaging. For each cell, 500 images were acquired with a cooled, electron-multiplying charge-coupled device camera (iXon DU-897; Andor) with an exposure time of 200 ms. For experiments with ZAP70 tSH2-GFP binding to monophosphorylated peptides 1.4 mW 488 nm light and an exposure time of 50 ms was used.

Trackmate software (62) was used to identify and track single molecules of ZAP70-Halotag-Alexa647 or ZAP70 tSH2-GFP. Spot intensities showed a single gausian distribution (Fig S13) indicating that single molecules were being tracked.

For peptide binding experiments a peptide surface was generated by coating a coverslip surface with poly-L-lysine-PEG-biotin (PLL(20)-g[3.5]- PEG(2)/PEG(3.4)- biotin(20%), SuSoS Surface Technology), saturating the biotin with 10 *µ*g/ml streptavidin in PBS/BSA (PBS with 1% w/v bovine serum albumin) for 30 min at room temp and incubating with biotinylated peptide. The surfaces were then washed thoroughly in PBS and ZAP70 tSH2-GFP or ZAP70 tSH2-Halo-Alexa647 was added at 200 nM in PBS/BSA with 1 mM DTT (to keep spurious disulfide bonds from forming) and the coverslips imaged as above for live cells.

The brightness and photobleaching of individual GFP molecules were determined by coating wells of a chamber coverslip in poly-L-lysine-PEG-biotin/streptavidin as above and adding a increasingly dilute biotinylated Avitag-GFP protein (produced in E.coli). For Alexa647, wells of a chamber coverslip were coated in poly-L-lysine and an increasing dilution of Alexa647-succinimidyl ester (Thermo Fischer) added and left to react for 30 min at room temperature. Wells with dilutions of Avitag-GFP or Alexa647 resulting in sparse single molecules with single step-bleaching characteristics were imaged.

### Analysis of particle tracking

Cumulative distributions of track survival probability vs time were fit with exponential decay models using Graphpad Prism software. One- and two-phase exponential fits were tested and an extra sum-of-squares F test was used to determine whether a two-phase decay was necessary. A single-phase decay was sufficient to fit experiments using monophosphorylated ITAM immobilised on glass coverslips resulting in unbinding rates that agreed with SPR experiments (Fig. S11). For data with immobilised biphosphorylated ITAM a one-phase decay did not fit well and two-phase decay was necessary. The slower phase component of this fit agreed well with SPR data. Similarly a one-phase decay did not fit data from live cell experiments well and a two-phase decay was used. The fast phase component of fits from bivalent peptide on glass and cellular data was similar (Fig. S12A,B) suggesting that it was a result of fluorophore photophysics due to fluorophore ‘blinking’. To demonstrate this, and to ensure that we were not missing an important short-lived ZAP70 state in T cells, we fixed T cells after stimulations and performed SPT finding that again, a two phase decay was needed with the slow phase now even slower than the ZAP70 SPR value (Fig. 5F, since it is fixed to the TIRF plane and tracks are lost by bleaching) but with a fast phase that again was similar to those observed before (Fig. S12A).

### Mathematical models

#### Operational model of kinetic proofreading (Fig. 1)

The system of ordinary-differential-equations (ODEs) describing the operational model of kinetic proof-reading with regulated unbinding (Fig. 1B) is as follows,

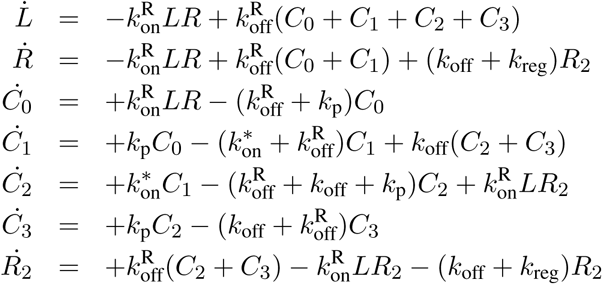

where all chemical states and reaction rates are depicted in Fig. 1B and parameter values summarised in Table 2. The initial conditions were *L*(*t* = 0) = *L*_*T*_, *R*(*t* = 0) = *R*_*T*_, and all other states 0. The model was integrated to steady state using the Matlab (Mathworks, MA) function *ode15s* for the indicated values of *k*_off_ and *k*_reg_. Specificity was defined as the ratio of *C*_3_ for the higher affinity agonist over *C*_3_ for the lower affinity non-agonist pMHC. Sensitivity was defined as the concentration of *C*_3_ for the higher affinity agonist normalised to the value obtained for a 0-step proofreading model. To calculate specificity and sensitivity using the standard kinetic proofreaidng model, we used the previously reported analytical solution (19, 60) with N=0, 1, 2, or 3 steps and all other parameters as indicated in Table 2.

**Table 2:** Parameters for operational model of kinetic proofreading (Fig. 1)

| Parameter | Description | Value | Units |
| --- | --- | --- | --- |
| $R_T$ | TCR concentration | 100 | molecules/$\mu\text{m}^2$ |
| $L_T$ | pMHC concentration | 1000 | molecules/$\mu\text{m}^2$ |
| $k_{\text{on}}^R$ | TCR/pMHC on-rate | 0.01 | $\mu\text{m}^2\text{s}^{-1}$ |
| $k_{\text{off}}^R$ | TCR/pMHC off-rate | $10^{-0.5}$ (agonist)<br>$10^{1.5}$ (non-agonist) | $\text{s}^{-1}$ |
| $k_p$ | Phosphorylation rate | 1 | $\text{s}^{-1}$ |
| $k_{\text{on}}^*$ | Effective first-order ZAP70 on-rate | 1 | $\text{s}^{-1}$ |
| $k_{\text{off}}$ | ZAP70 off-rate | variable | $\text{s}^{-1}$ |
| $k_{\text{reg}}$ | Regulated ZAP70 off-rate | variable | $\text{s}^{-1}$ |

### Bivalent ZAP70/ITAM binding model (Fig. 2)

The bivalent interaction of ZAP70 (with tandem SH2 domains) binding to phosphorylated ITAMs (with two phosphorylated tyrosines) that can be dephosphorylated by a phosphatase (Fig. 2D) is modelled using the following ordinary-differential-equations (ODEs),

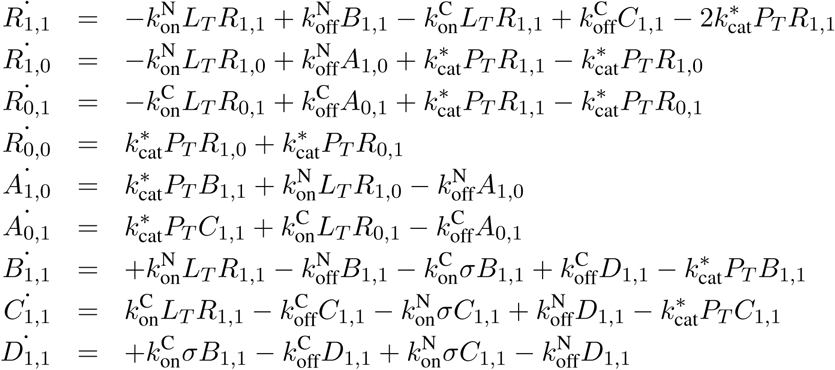

where subscripts indicate which of the two tyrosines are phosphorylated (e.g. 1,0 and 1,1 indicate phosphorylation of ITAM N only or both ITAM N and C, respectively), *R* represents unbound ITAM states with the indicated phosphorylation, *A* represents bound ITAM states that are monophosphorylated, *B* and *C* are biphosphorylated ITAM states where only ITAM N or ITAM C are bound by a ZAP70 SH2 domain, respectively, and *D* is a biphosphorylated ITAM with both tyrosines bound to the tandem SH2 domains of ZAP70. The model parameters are described in the text and summarised in Table 3. The model was numerically solved in Matlab (Mathworks, MA) using *ode15s*.

**Table 3:**
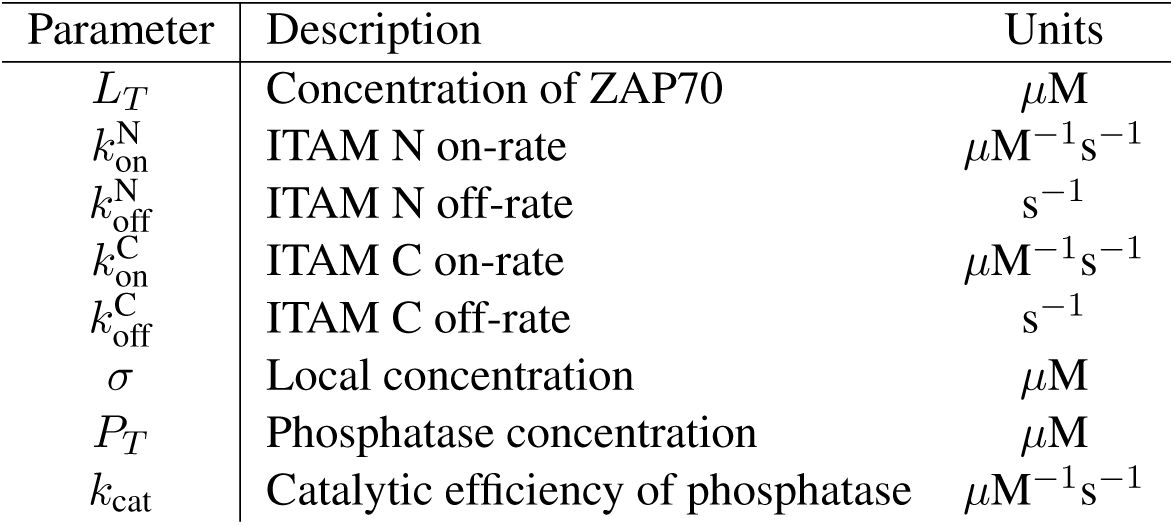
Parameters for bivalent ZAP70/ITAM binding model

The mathematical model makes several assumptions that we make explicit here. First, phosphatases can only dephosphorylate tyrosines that are not bound to SH2 domains, as previously described (31). Second, the absolute amount of phosphorylated tyrosines is small compared to the absolute amount of ZAP70 or phosphatases so that they are not depleted, which allows for these molecules to have a fixed concentration (*L*_*T*_ and *P*_*T*_, respectively). This is a standard assumption when modelling SPR experiments where they are replenished by flow and is reasonable for cellular experiments because only a fraction of TCRs will be phosphorylated. Third, only a single molecule of ZAP70 can interact with an ITAM. This is reasonable because the maximum concentration of ZAP70 binding to ITAM3 NC (0.6 *µ*M) is much lower than the experimentally observed *σ* (1161 *µ*M) so that solution ZAP70 could not compete with ZAP70 already bound to the ITAM. Fourth, no cross reactions were included that would allow, for example, ITAM N to bind not only C-SH2 of ZAP70 but also the N-SH2. This is reasonable because we could not find any evidence for these reactions and this is underlined by the fact that the present simpler model was sufficient to explain the data. Fifth, ZAP70 binding is defined as all states where ZAP70 is interacting with an ITAM (i.e. *A*_1,0_ + *A*_0,1_ + *B*_1,1_ + *C*_1,1_ + *D*_1,1_). This is reasonable because both in SPR and in microscopy, the microscopic states of ZAP70 are not resolved but rather total localisation is reported.

The calculations in Fig. 2 were initialised with all states 0 except *R*_1,1_ which was set to 1. The concentration of ZAP70 remained fixed for the entire calculation (*L*_*T*_ = 1 *µ*M) whereas the concentration of phosphatase (*P*_*T*_) was initially 0 for 30 seconds to allow the system to come to steady-state before increasing to the indicated concentration with *k*_cat_ = 0.1 *µ*M^−1^s^−1^. The bivalent calculations (Fig. 2D,E) used 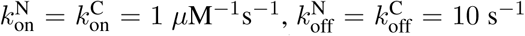, and *σ* = 1000 *µ*M resulting in an effective unbinding rate back to solution of 0.2 s^−1^. The monovalent calculations (Fig. 2A,B) used the bivalent model with 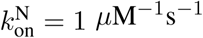 and 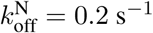 but with 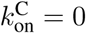 so that only a single SH2 domain can interact.

The SPR experiments provide the amount of ZAP70 bound at steady-state for different concentrations of ZAP70 on the biphosphorylated ITAM NC peptide. To calculate this in the mathematical model above, we set *P*_*T*_ = 0 (no phosphatase) and solved the ODE system in the steady-state to calculate total ZAP70 binding (*Y*, defined as *Y* = *A*_1,0_ + *A*_0,1_ + *B*_1,1_ + *C*_1,1_ + *D*_1,1_),

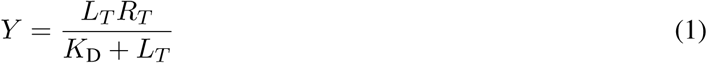

where *R*_*T*_ is the total peptide concentration, *L*_*T*_ is the ZAP70 concentration, and *K*_D_ is the effective ZAP70 affinity 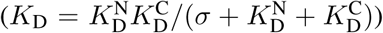. This equation derived from the bivalent model has an identical form to that of the monomeric binding, which explains why an apparent monovalent model is able to fit the bivalent interaction of ZAP70.

#### Molecular model of kinetic proofreading (Fig. S9)

The rule-based framework of BioNetGen (66) was used to generate a molecular model of ZAP70 activation (see Supplementary Information for the full code). This framework is particularly suited for molecular models where the number of chemical states can become very large as a result of reactions that take place independently. The model included TCR/pMHC binding that initiated ITAM phosphorylation (2 sites), ZAP70 binding by kinetic avidity (using the experimentally determined parameters), and ZAP70 phosphorylatio on 4 sites. Kinetic proofreading was realised by allowing for phosphorylation (ITAM and ZAP70) only when the TCR is bound and dephosphorylation only when the TCR is unbound. Importantly, ZAP70 binding was dependent on ITAM phosphorylation but independent of TCR/pMHC binding. In this way, regulated unbinding of ZAP70 naturally arose out of the explicit modelling of kinetic avidity and phosphatase activity. These reactions produced a total of 64 distinct chemical states and 270 chemical reactions between these states (see Fig. S9 for a schematic of a subset of these states and reactions).

The model was numerically solved in Matlab (Mathworks, MA) using *ode15s* to the steady-state. The amount of TCR bound ZAP70 (either with one or two SH2 domains) and the amount of activated ZAP70 (defined as the amount of bound ZAP70 that is phosphorylated on all 4 sites) was calculated for different TCR/pMHC off-rates using different phosphatase activities at the membrane. The model parameters are summarised in Table S4. To model ZAP70 binding by kinetic avidity the experimental parameters for each individual SH2 domain were used (Table 1) along with the estimate of the local concentration (*σ* = 1161 *µ*M). To model ZAP70 binding by static avidity, kon2 in the model was set to zero with kon1 and koff1 set equal to the rates determined for ZAP70 binding ITAM NC (Table 1).

**Table 4:** Parameters for molecular model of kinetic proofreading

| Parameter | Description | Value | Units |
| --- | --- | --- | --- |
| $R$ | Number of TCRs | 31000 | - |
| $L$ | Number of pMHCs | 310000 | - |
| $Z$ | Number of ZAP70s | 130000 | - |
| $k_{\text{on}}$ | TCR/pMHC on-rate | 0.01 | $\mu\text{m}^2/\text{s}$ |
| $k_{\text{off}}$ | TCR/pMHC off-rate | varied | $\text{s}^{-1}$ |
| $k_{\text{pr}}$ | Phosphorylation rate of ITAM | 1 | $\text{s}^{-1}$ |
| $k_{\text{pe}}$ | Phosphorylation rate of ZAP70 | 4 | $\text{s}^{-1}$ |
| $k_{\text{m}}$ | Dephosphorylation rate at the membrane | varied | $\text{s}^{-1}$ |
| $k_{\text{c}}$ | Dephosphorylation rate in cytosol | $10^8$ | $\text{s}^{-1}$ |

#### Stochastic spatial simulations (Fig. S7)

To investigate the rebinding of a cytosolic protein (particle) to a surface receptor, we utilised two independent spatial stochastic simulation algorithms that produced identical results. The Brownian Dynamics algorithm (63) is a continuous-space discrete-time algorithm whereas the Gillespie algorithm is a discrete-space (on lattice) continuous-time algorithm (64, 65). The surface receptor was assumed to be fixed (immobile) at the origin (*r* = 0). The cytosolic protein was assumed to diffuse in the cytoplasm (*D* = 10 *µ*m^2^/s) with diffusion is limited to the half-space below the membrane (i.e. reflecting boundary at the membrane). The particle can reversibly bind to the receptor with first-order kinetics (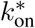 and *k*_off_ = 0.2 s^−1^), provided it is within a distance *b* = 10 nm of the origin, where *b* is the distance explored by the cytoplasmic tail of the receptor. The on-rate was proportional to the bimolecular on-rate, the number of binding sites, and the volume (proportional to b): 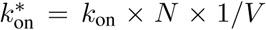, where *k*_on_ = 1.89 *µ*M^−1^s^−1^, *N* is the number of ITAMs, and V is the reaction volume (*V* = ((4*/*3)*πb*^3^)*/*2). The simulations were initialised with the cytosolic protein having just unbound from the receptor and therefore, it was placed randomly within the half-sphere determined by *b*. The simulations were terminated when the cytosolic protein diffused beyond *r* = *a*. We took *a* = 150 nm, which is approximately the mean distance between ZAP70 molecules within the cytoplasm. The simulations were repeated 10^6^ times and the mean first passage time to reach *a* and the number of rebinding events within this time were recorded for different values of *N* .

## Supplementary Information

### Supplementary Figures

**Figure S1:**
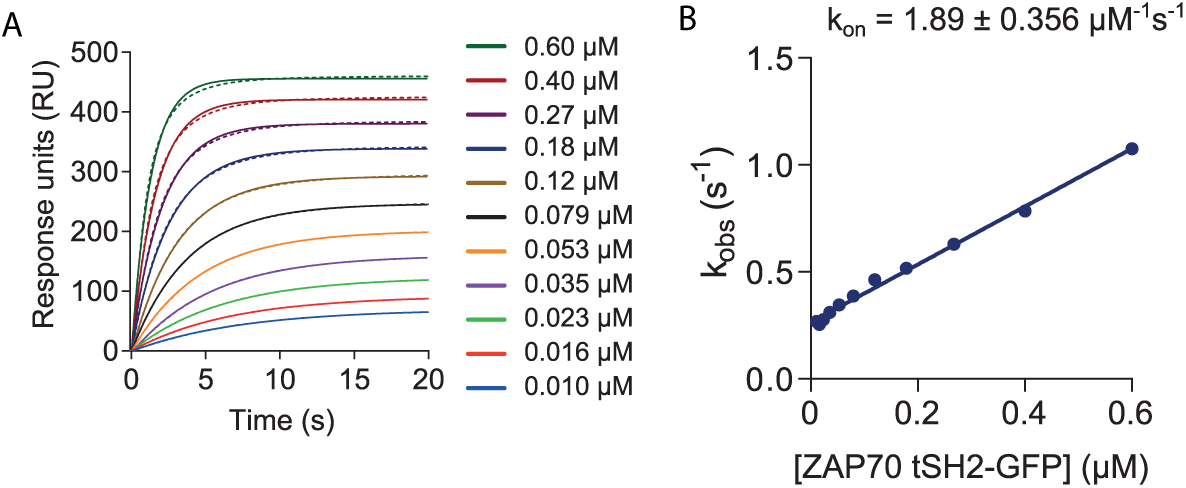
k_obs_ analysis of tSH2-GFP interacting with biphosphorylated ITAM3. (A) Representative association phase data (dots) and exponential association phase fit (solid lines). (B) Fitted rate constant (k_obs_) over [tSH2-GFP] reveals the expected linear relationship (k_obs_ = *k*_on_[tSH2-GFP]+*k*_off_). The value of *k*_on_ (slope) is indicated. Related to Fig. 3

**Figure S2:**
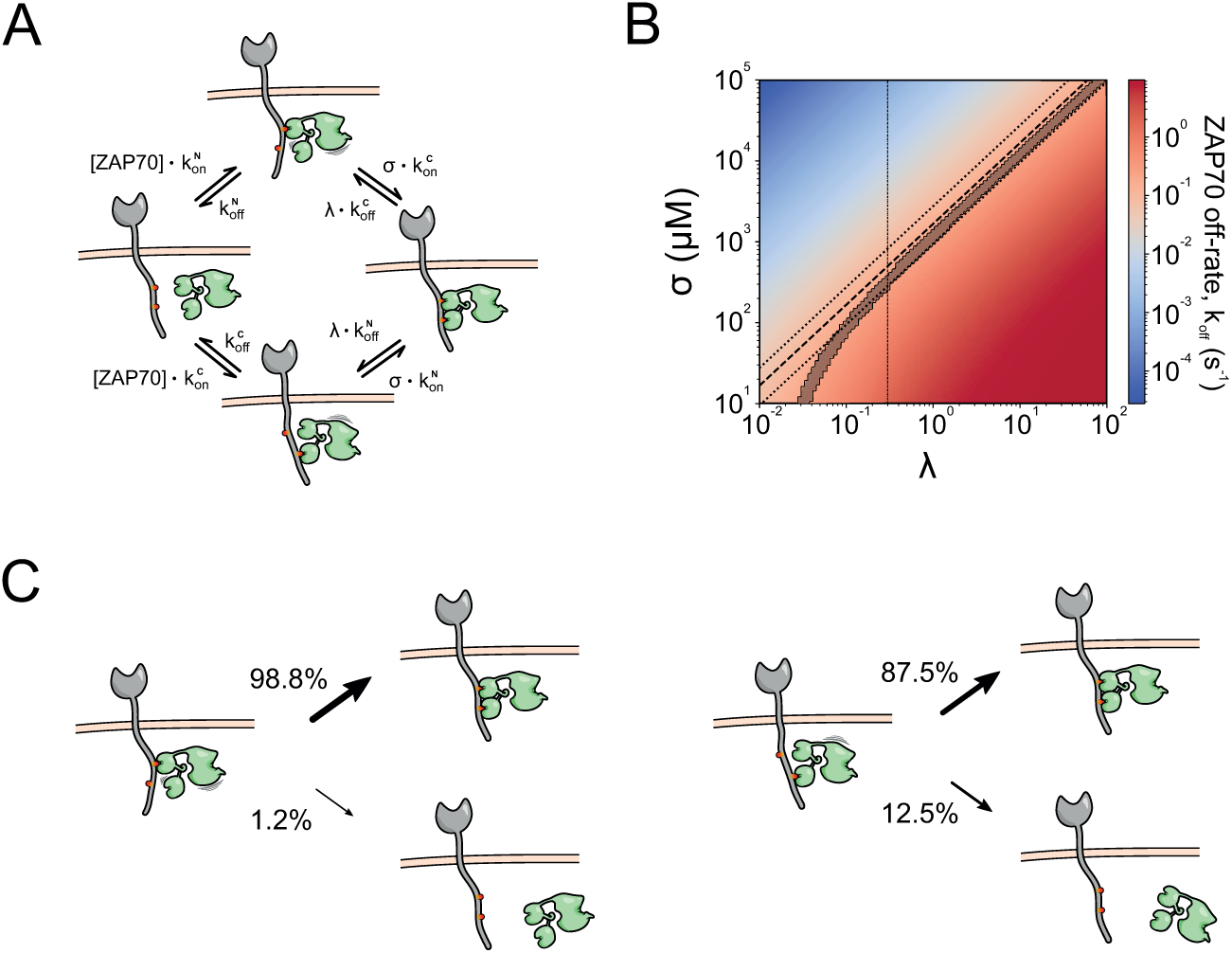
Model of ZAP70/ITAM interaction that includes conformational cooperativity predicts kinetic avidity. (A) The model of ZAP70/ITAM interaction (Fig. 2D) was modified to include a conformational cooperativity parameter (*λ*) that multiplied the off-rates only when ZAP70 is bound to both ITAM tyrosines. Since *λ* modifies the 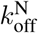 and 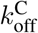 rates, values of *λ <* 1 represent positive co-operativity, whereas *λ >* 1 represent negative co-operativity. (B) Heap map of ZAP70 off-rate predicted by the mathematical model when varying *σ* and *λ*. As before, the 4 kinetic rate constants were fixed to their experimentally determined values (Table 1). Regions within the range of *k*_off_ ±SEM measured experimentally are shaded in grey. The dashed line on the diagonal is the experimentally determined ratio of *σ/λ* using the dissociation constants 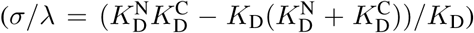 with the dotted lines indicating upper and lower bounds. The best overlap occurs for values of *λ >* 0.3 (indicated by a dashed vertical line). (C) Probability of rebinding or unbinding when ZAP70 is bound to N- (left panel) or C-terminal (right panel) tyrosines of ITAM3 if *λ* = 0.3 and *σ* = 300 *µ*M.

**Figure S3:**
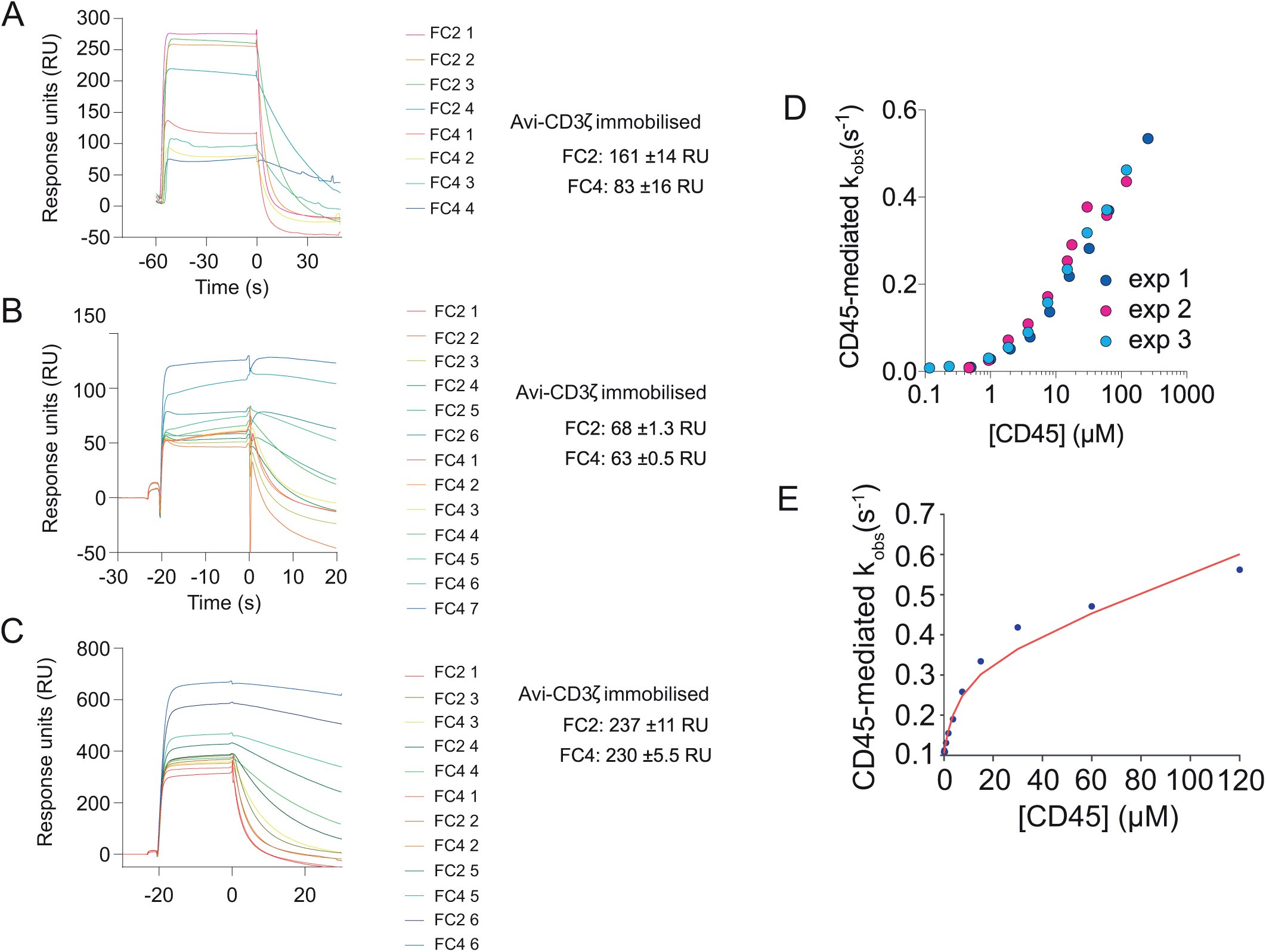
The CD45-mediated observed off-rate of ZAP70 is similar at different Avi-CD3*ζ* ITAM3 peptide immobilisation levels. (A-C) Overlayed sensograms from individual experiments differing in Avi-CD3*ζ* ITAM3 peptide immobilisation. The mean ±SD of peptide immobilised per cycle is shown to the right of the sensograms. (D) The fitted CD45-mediated observed off-rate over [CD45] for experimental data shown in panels A (exp 1), B (exp 2) and C (exp 3). (E) Catalytic efficiecy of CD45 dephosphorylating CD3*ζ* ITAM3. Given that all ZAP70 binding kinetics were identified and that the concentration of CD45 is known, we were able to directly fit the CD45-mediated *k*_obs_ (red curve) to estimate the catalytic efficiecny (*k*_cat_*/K*_M_) of CD45 dephosphorylating ITAM3 to be 0.103±0.01 *µ*M^−1^ s^−1^. Representative data and fit are shown (N=4).

**Figure S4:**
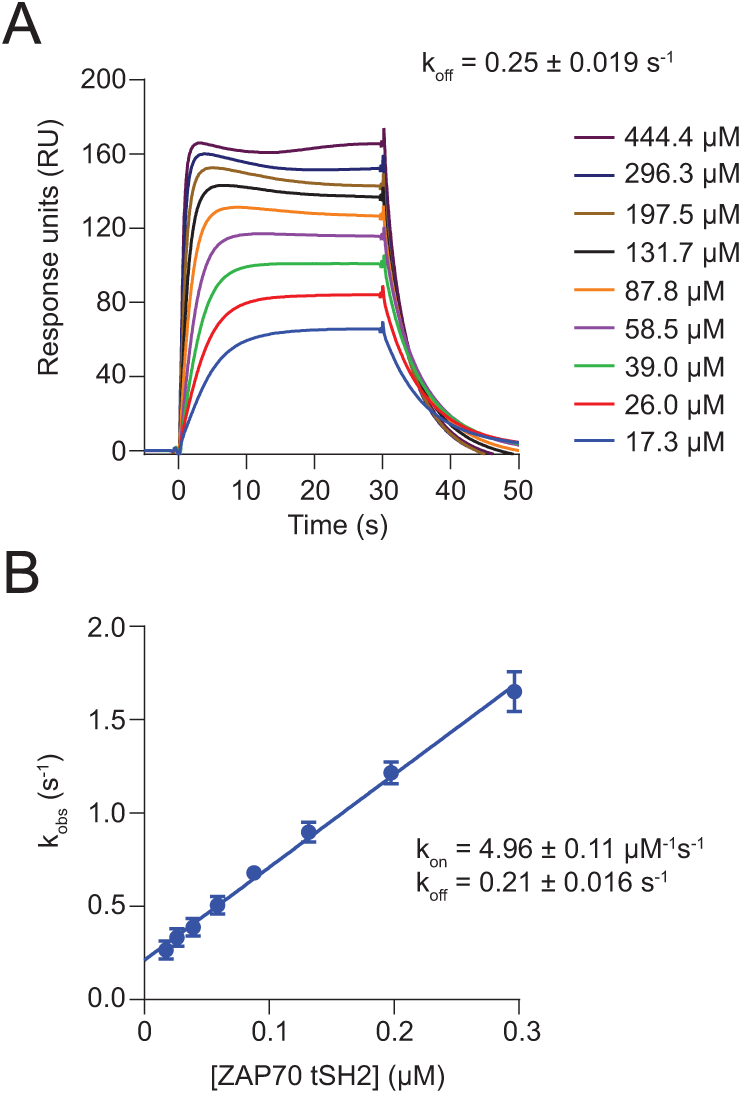
Binding kinetics of ZAP70 interacting with full length intracellular CD3*ζ* peptide phosphorylated on ITAM3. (A) Example sensograms from ZAP70 tSH2-GFP protein binding to phosphorylated Avi-CD3*ζ*. The dissociation rate (*k*_off_) derived from fits of the dissociation phase of 4 experiments ±SEM is shown. (B) Plot of k_obs_ vs concentration of tSH2-GFP, *k*_on_ and *k*_off_ ±SE from the slope and intercept of the linear regression are shown.

**Figure S5:**
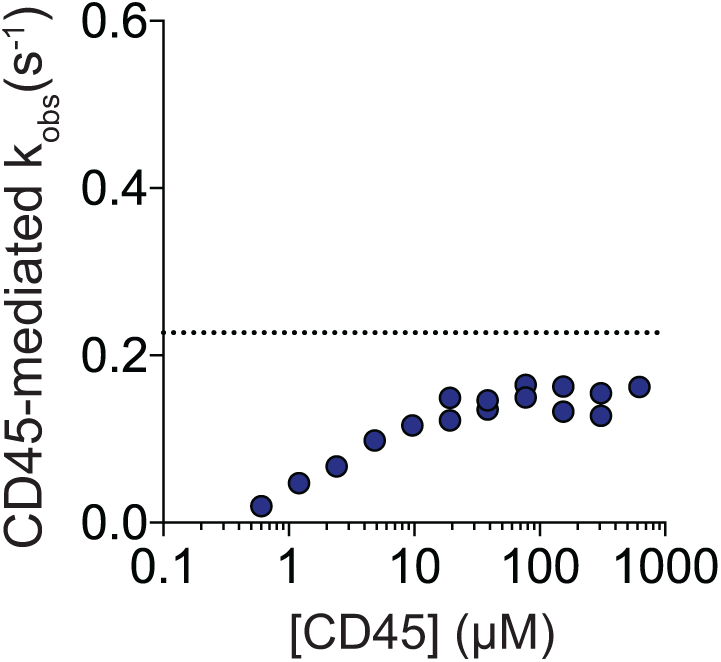
The CD45-mediated observed off-rate of ZAP70 tSH2-GFP measured on short biphosphorylated ITAM3 peptide over [CD45]. In contrast to data for the full-length intracellular domain (shown in Fig. 4), data with this shorter peptide shows that the observed off-rate plateaus near the value of *k*_off_ for ZAP70 tSH2-GFP for ITAM3 (shown as a horizontal dotted line).

**Figure S6:**
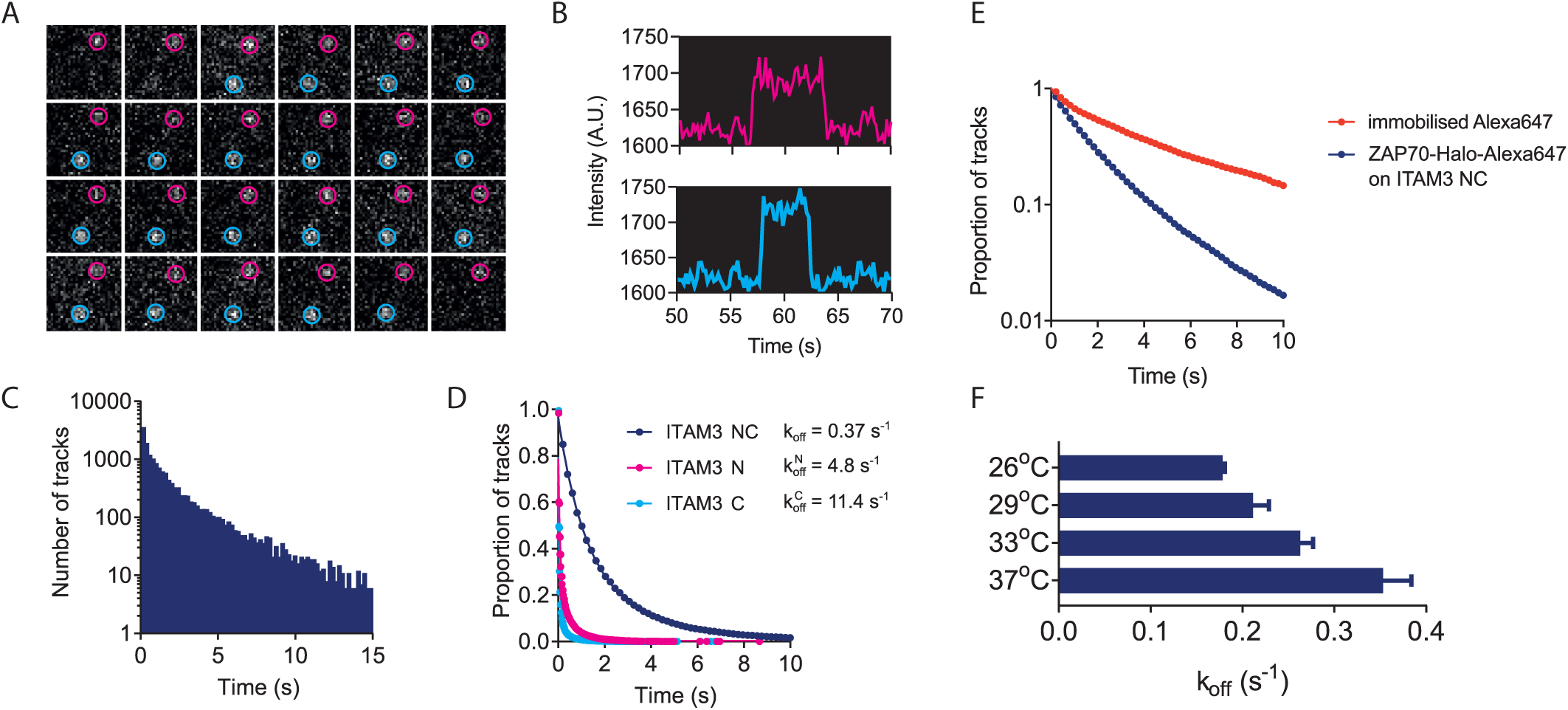
Unbinding kinetics measured by SPT using TIRF microscopy are similar to those measured by SPR. (A) Example 2.9 x 2.9 *µ*m region from TIRF imaging showing single ZAP70 tSH2-GFP molecules (outlined in magenta and teal circles) binding to biphosphorylated CD3*ζ* ITAM3 peptide immobilised on a glass coverslip. Images were acquired at 5 Hz. (B) Intensity vs time trace of regions containing particles in panel A showing single step increases and decreases indicative of single molecule binding and unbinding events. (C) Example histogram of track lengths from SPT of ZAP70 tSH2-GFP binding to biphosphorylated CD3*ζ* ITAM3 peptide immobilised on a coverslip surface. Images were acquired at 5 Hz for 400 seconds. (D) Track lifetimes of tSH2-GFP binding to N-terminally phosphorylated, C-terminally phosphorylated and biphosphorylated CD3*ζ* ITAM3 peptide with eponential fit (line). Data are representative of 16 image stacks taken over 3 independent experiments. (E) Photobleaching minimally impacts on *k*_off_ measurements. Bleaching rate of Alexa647-PLL sparsely immobilised on glass (immobilised Alexa647) compared with the lifetime of ZAP70 tSH2-Halotag labelled with Alexa647-haloligand binding to immobilised biphosphorylated ITAM3 peptide (ZAP70-Halo-Alexa647 on ITAM3 NC). Both were imaged at 37°C using identical imaging conditions. (F) *k*_off_ of ZAP70 tSH2-GFP binding to biphosphorylated ITAM3 peptide at different temperatures demonstrating the ability to measure longer binding lifetimes.

**Figure S7:**
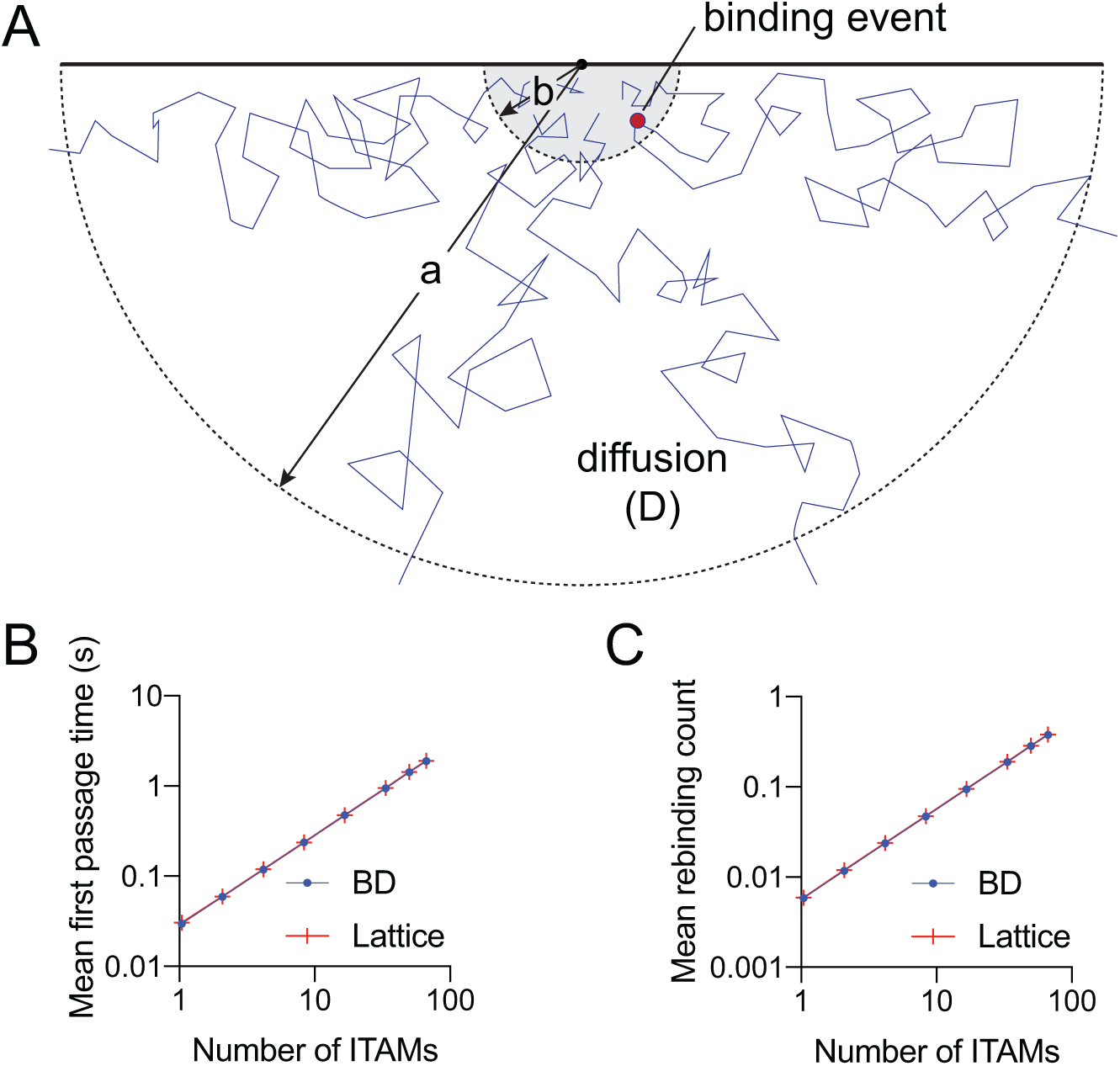
Stocahstic spatial simulations reveal limited rebinding between the same cytosolic protein and the same surface receptor. A) Simulation schematic. The cytoplasmic tail of the surface receptor is assumed to explore the half-sphere of radius *b* and the cytosolic protein can rebind the receptor within this half-sphere with first-order kinetics. The simulation is initiated with the cytosolic protein unbound from the surface receptor and placed randomly within the binding half-sphere. The simulation is terminated when the cytosolic protein diffuses beyond the half-sphere of radius *a*. B) The average time to diffuse beyond the half-sphere of radius *a* (mean first passage time) over the number of binding sites. C) The average number of rebinding events over the number of binding sites. See Methods for simulation details.

**Figure S8:**
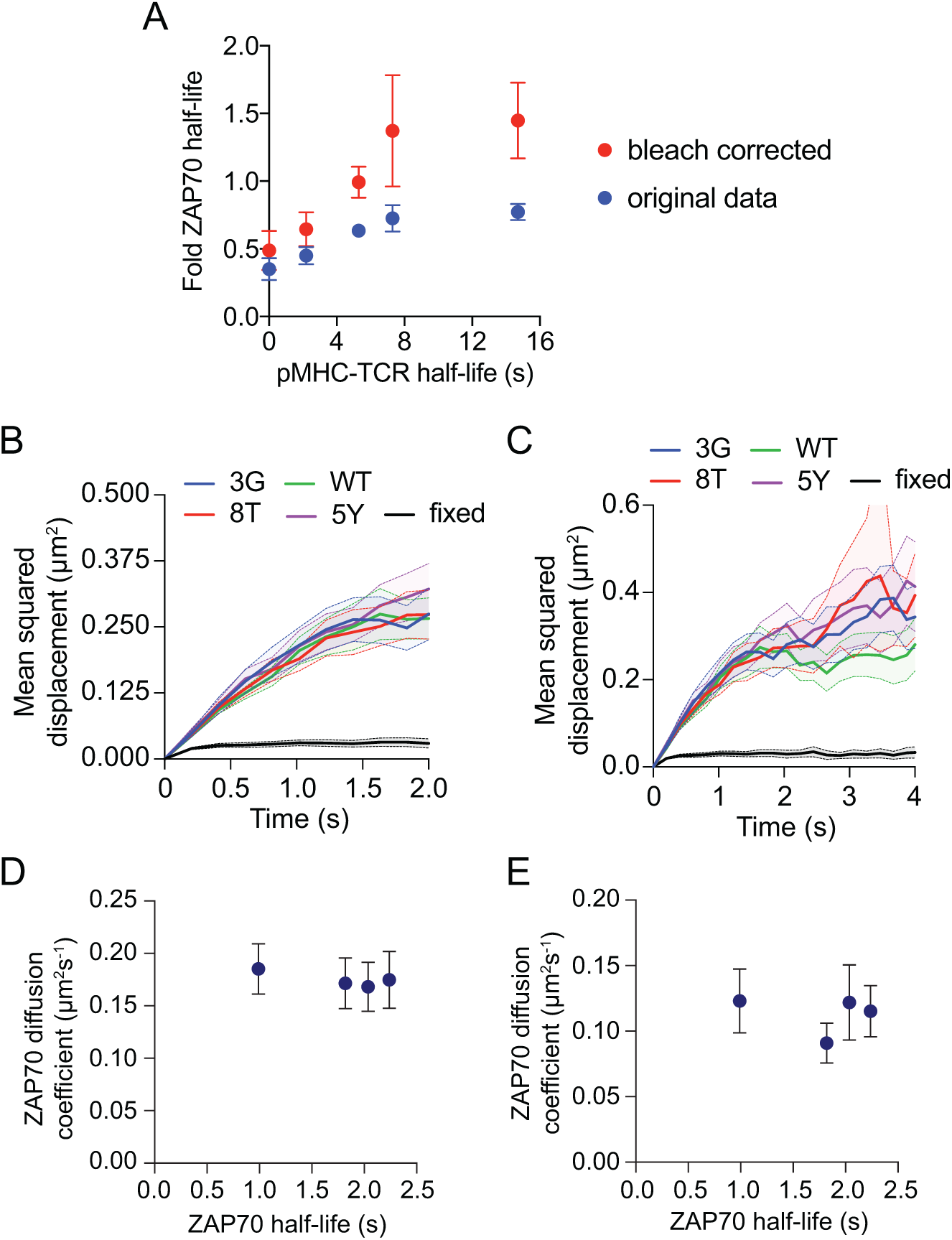
No evidence that the same ZAP70 protein rebinds the TCR. A) The ZAP70 membrane half-life does not exceed the SPR half-life for all TCR/pMHC interactions tested. B-C) The mean squared displacement is similar on short (panel B) and long (panel C) timescales for different pMHCs. D-E) Diffusion coefficients derived from the truncated short timescale data (panel D) and the complete data (panel E) do not correlate with the ZAP70 membrane half-life .

**Figure S9:**
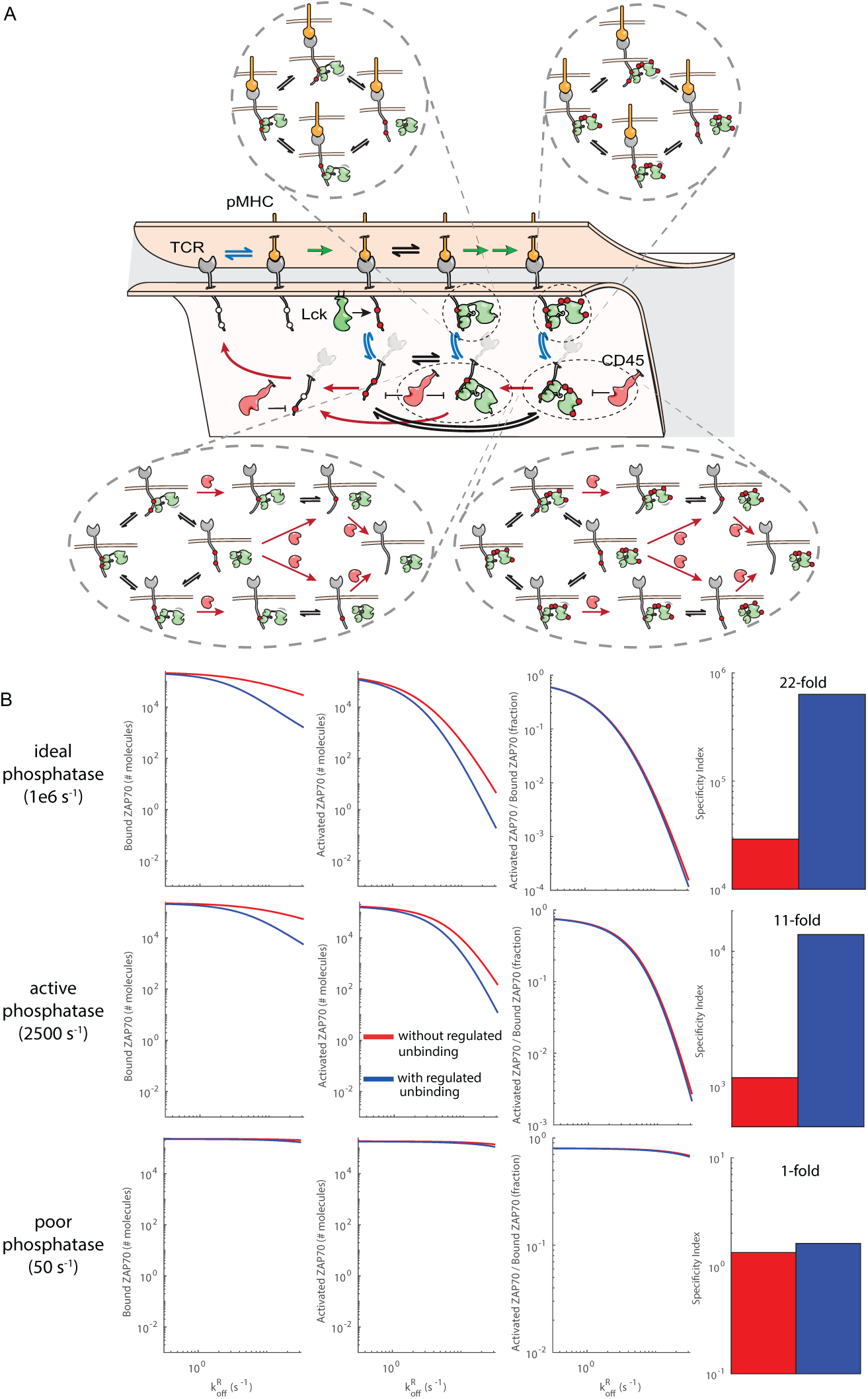
Molecular model of kinetic proofreading. A) Simplified schematic of the molecular proofreading model that includes TCR/pMHC binding (blue arrows), phosphorylation of the TCR and ZAP70 when the TCR is bound to pMHC (green arrows), dephosphorylation when the TCR is unbound from pMHC (red arrows), and ZAP70 binding (black arrows). Bound ZAP70 is defined by the amount of ZAP70 interacting with the TCR with either one or two SH2 domains and activated ZAP70 is defined as the amount of bound ZAP70 fully phosphorylated. B) The amount of ZAP70 bound (1st column), activated (2nd column), or the ratio of activated to bound (3rd column) over the TCR/pMHC off-rate (x-axis) when using an ideal (first row), an active (second row), or a poor (third row) phosphatase. As in Fig. 1, specificity (4th column) was defined as the ratio of active ZAP70 between an agonist 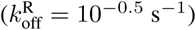 and non-agonist 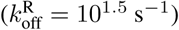 pMHC. Specificity is reduced with a poor phosphatase because sustained signalling abolishes proofreading and consistent with the operational model, ZAP70 binding by kinetic avidity exhibits higher specificity than static avidity because unbinding can be regulated. Although appreciable ZAP70 binding is predicted at all TCR/pMHC off-rates, the amount of activated ZAP70 is nearly ∼1000-fold lower for large off-rates compared to small off-rates. See Methods for computational details.

**Figure S10:**
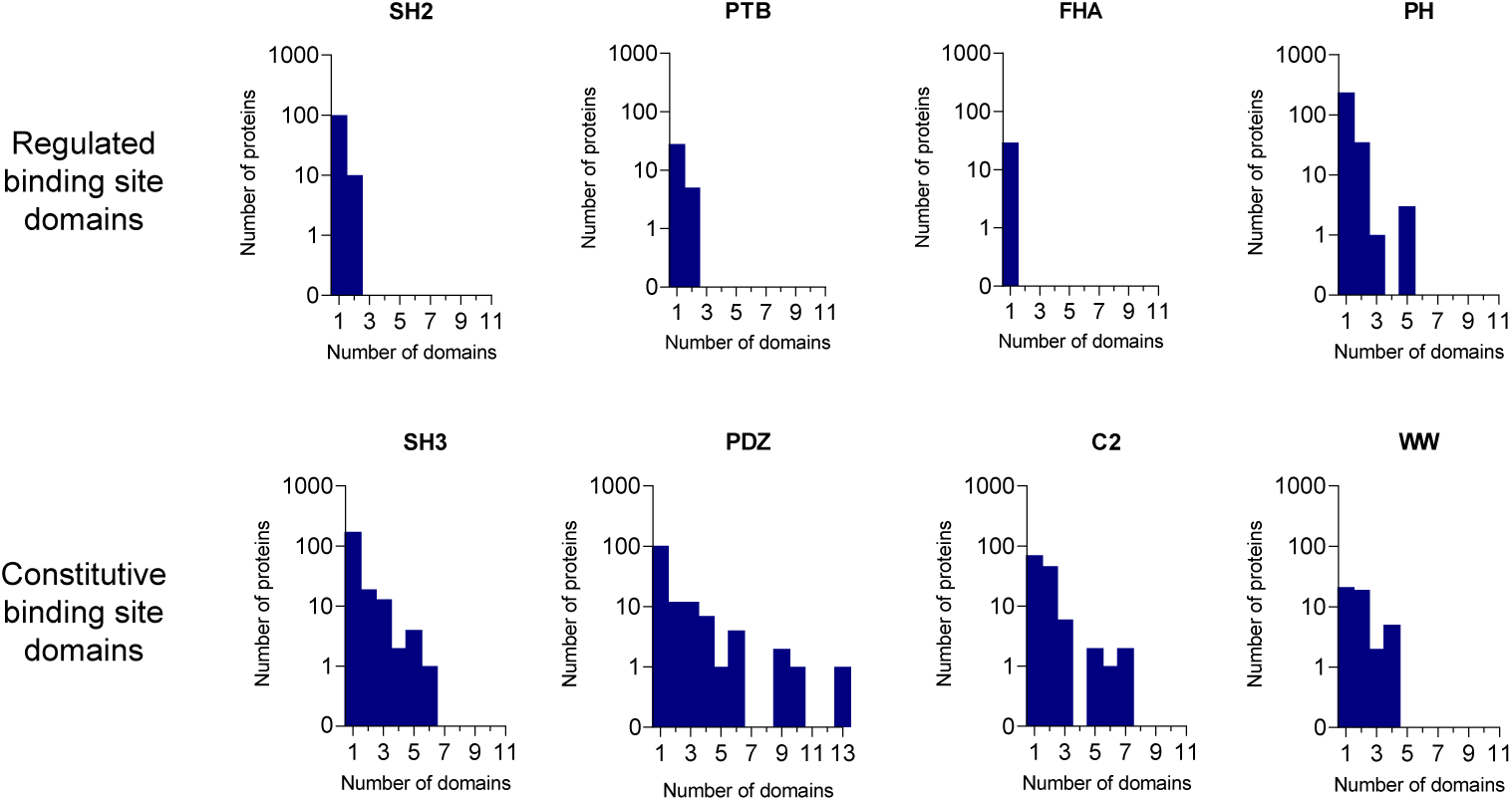
Numbers of genes in the human genome with single or multiple repeated binding domains. Data for Src homology 2 (SH2), Src homology 3 (SH3), C2, Pleckstrin homology (PH), phosphotyrosine binding (PTB), PDZ, forkhead-associated (FHA) and WW domains are shown. Shown are the number of proteins (y-axis) that have the indicated number of binding domains (x-axis) for each domain family. Note that SH2, PTB, FHA, and PH domains bind to regulated sites that can be either ‘on’ or ‘off’ (e.g. SH2 and PTB domains bind to phosphorylated but not dephosphorylated tyrosines) whereas SH3, PDZ, C2, and WW domains bind to sites that are constitutively ‘on’. Domains included in the survey are the most well-characterized domains and all data is obtained from the SMART domain database.

**Figure S11:**
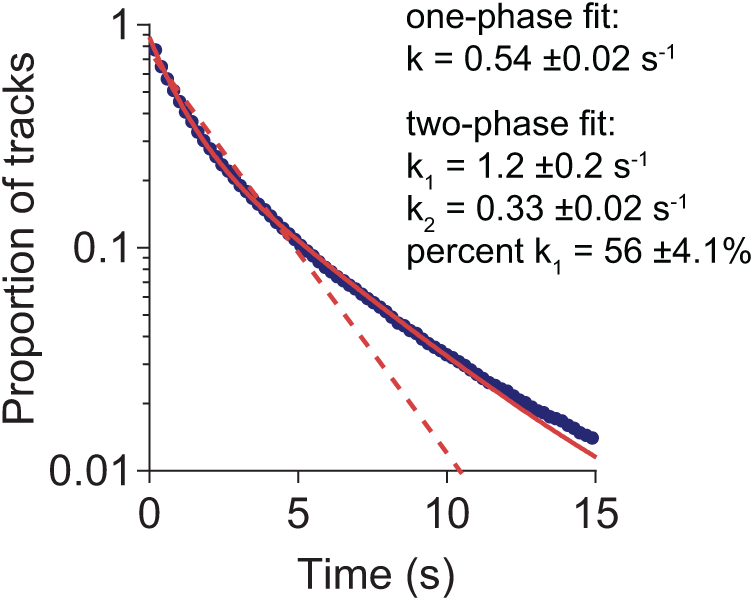
One- and two-phase exponential decay fits of ZAP70 tSH2-Halo Alexa647 binding to bivalent ITAM3. A two-phase decay model (solid red line) fit the track lifetime data (navy blue dots) significantly better than a one-phase decay (dashed red line). Average decay rates ± SEM are shown (n=9).

**Figure S12:**
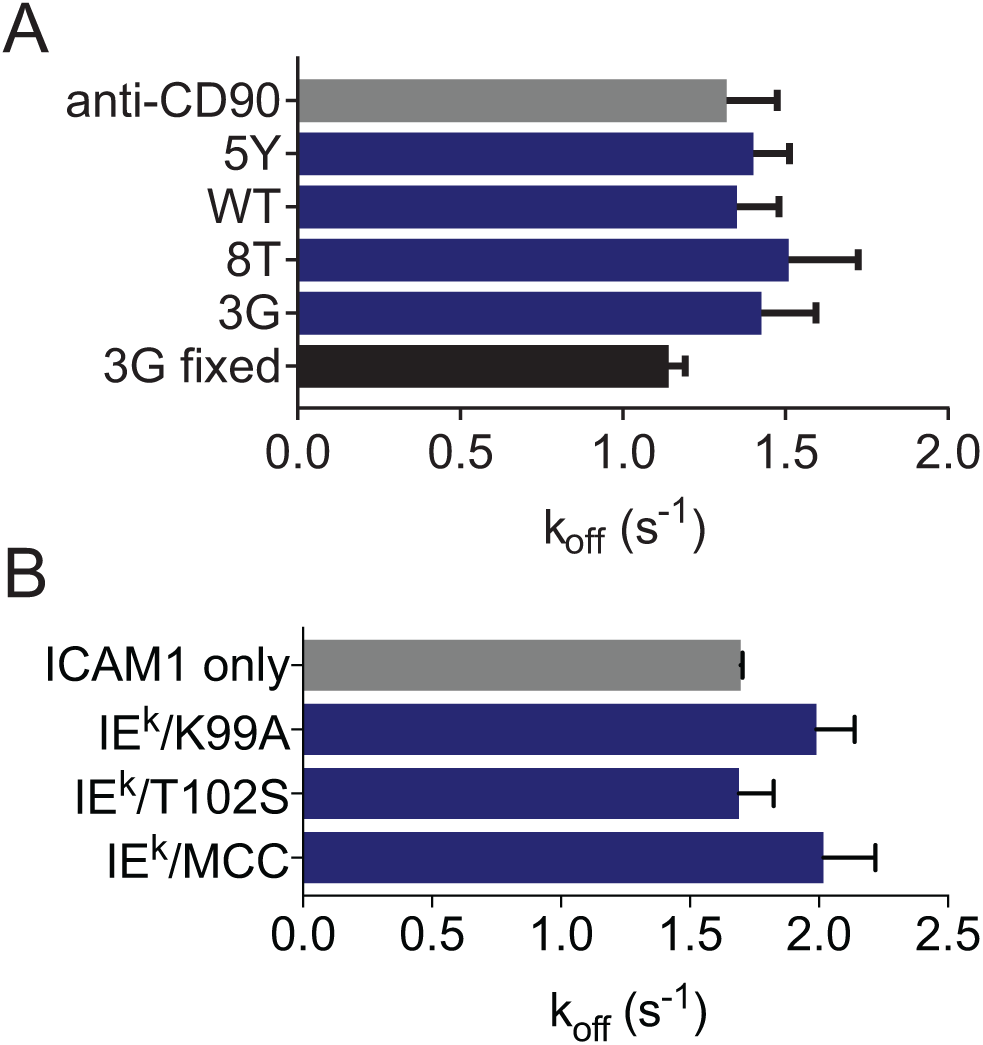
Fast phase decay rate component of fits from live cells SPT experiments. (A) Fast decay rates from fits of ZAP70 Halo-Alexa647 SPT data in ILA Jurkats stimulated with the indicated pMHC in live cells stimulated with the indicated pMHC or control (anti-CD90) or cells fixed after 5 min stimulation with 3G pMHC (3G fixed). Corresponding slow phase rates are shown in Fig 5E. (B) Fast decay rates from fits of ZAP70-GFP SPT data in primary mouse AND T cells stimulated with the indicated pMHC. Corresponding slow phase rates are shown in Fig 5I. Data are from at least 8 cells per condition imaged in three separate experiments. Mean ±SEM are shown. There were no significant differences between conditions (one-way ANOVA with Tukeys post-test).

**Figure S13:**
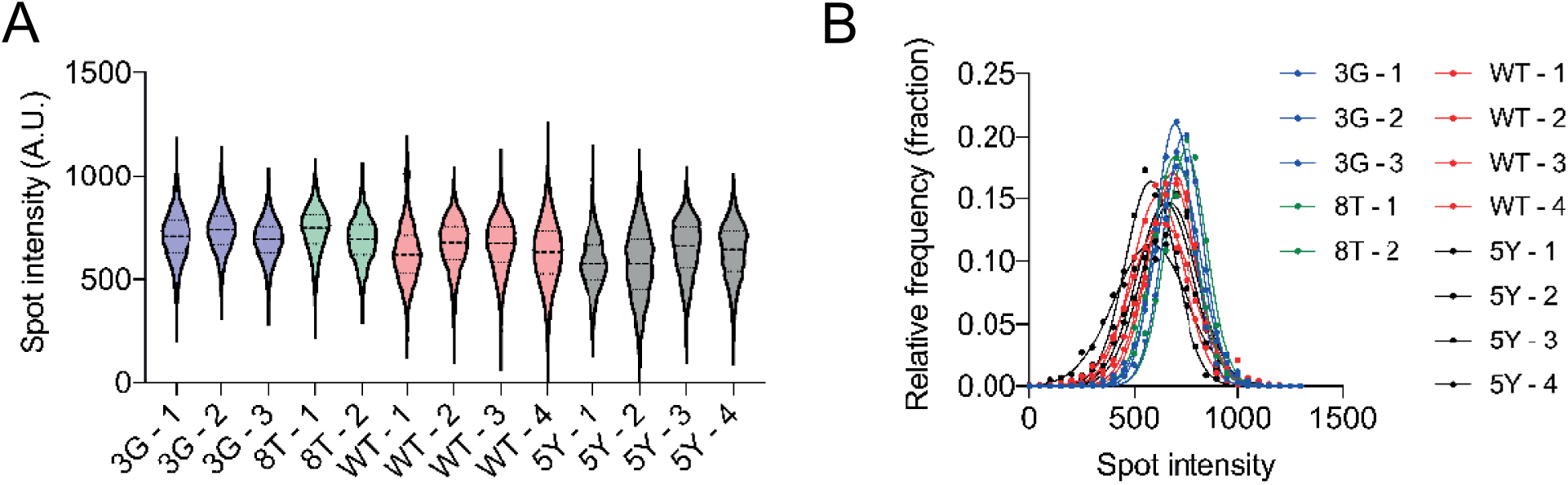
Spot intensities from single particle tracking. (A) Background subtracted spot intensities from image series of cells stimulated with the indicated pMHC. Data from individual cells in one representative experiment are shown. (B) Data in (A) plotted as overlapping histograms.

### BioNetGen BNGL file for molecular model of kinetic proofreading

~~~
#BNGL file for molecular model for kinetic proofreading

begin parameters

#TCR/pMHC binding (koff varied)
kon 0.01
koff 1

#ZAP70 Binding
kon1 0.26
koff1 5.5
kon2 1.52
koff2 11.4
sigma 1161

#ZAP70 Intramolecular reactions
kon1s   kon1*sigma
koff1s  koff1
kon2s   kon2*sigma
koff2s  koff2

#TCR Phosphorylation (when pMHC is bound to TCR)
kpr     1

#ZAP70 Phosphorylation (when pMHC and ZAP70 are bound to TCR)
kpe     4

#Dephosphorylation rate at the membrane (varied)
km      2500

#Dephosphorylation rate in solution
kc      1e8

end parameters

begin molecule types
    L(b)
    R(b,Y1∼U∼P,Y2∼U∼P)
    Z(b1,b2,Y∼U∼P∼2P∼3P∼4P)
end molecule types

begin seed species
    L(b)             310000
    R(b,Y1∼U,Y2∼U)   31000
    Z(b1,b2,Y∼U)     130000
end seed species

begin observables
   Molecules BoundReceptor R(b!+)
   Molecules BoundEffector Z(b1!+,b2),Z(b1,b2!+),Z(b1!+,b2!+)
   Molecules BoundActiveEffector Z(b1!+,b2,Y∼4P),Z(b1,b2!+,Y∼4P),Z(b1!+,b2!+,Y∼4P)
   Molecules ActiveEffector Z(Y∼4P)
end observables

begin reaction rules

#TCR/pMHC binding
L(b) + R(b) <-> L(b!1).R(b!1) kon,koff

#TCR phosphorylation
R(b!+,Y1∼U) -> R(b!+,Y1∼P) kpr
R(b!+,Y2∼U) -> R(b!+,Y2∼P) kpr

#ZAP70 binding
R(Y1∼P,Y2∼U) + Z(b1,b2) <-> R(Y1∼P!1,Y2∼U).Z(b1!1,b2) kon1,koff1
R(Y1∼U,Y2∼P) + Z(b1,b2) <-> R(Y1∼U,Y2∼P!1).Z(b1,b2!1) kon2,koff2

R(Y1∼P,Y2∼P) + Z(b1,b2) <-> R(Y1∼P!1,Y2∼P).Z(b1!1,b2) kon1,koff1
R(Y1∼P,Y2∼P) + Z(b1,b2) <-> R(Y1∼P,Y2∼P!1).Z(b1,b2!1) kon2,koff2

R(Y1∼P!1,Y2∼P).Z(b1!1,b2) <-> R(Y1∼P!1,Y2∼P!2).Z(b1!1,b2!2) kon2s,koff2s
R(Y1∼P,Y2∼P!1).Z(b1,b2!1) <-> R(Y1∼P!2,Y2∼P!1).Z(b1!2,b2!1) kon1s,koff1s

#ZAP70 phosphorylation
Z(b1!+,b2,Y∼U) -> Z(b1!+,b2,Y∼P) kpe
Z(b1!+,b2,Y∼P) -> Z(b1!+,b2,Y∼2P) kpe
Z(b1!+,b2,Y∼2P) -> Z(b1!+,b2,Y∼3P) kpe
Z(b1!+,b2,Y∼3P) -> Z(b1!+,b2,Y∼4P) kpe

Z(b2!+,b1,Y∼U) -> Z(b2!+,b1,Y∼P) kpe
Z(b2!+,b1,Y∼P) -> Z(b2!+,b1,Y∼2P) kpe
Z(b2!+,b1,Y∼2P) -> Z(b2!+,b1,Y∼3P) kpe
Z(b2!+,b1,Y∼3P) -> Z(b2!+,b1,Y∼4P) kpe

Z(b2!+,b1!+,Y∼U) -> Z(b2!+,b1!+,Y∼P) kpe
Z(b2!+,b1!+,Y∼P) -> Z(b2!+,b1!+,Y∼2P) kpe
Z(b2!+,b1!+,Y∼2P) -> Z(b2!+,b1!+,Y∼3P) kpe
Z(b2!+,b1!+,Y∼3P) -> Z(b2!+,b1!+,Y∼4P) kpe

#Receptor dephosphorylation
R(b,Y1∼P) -> R(b,Y1∼U) km
R(b,Y2∼P) -> R(b,Y2∼U) km

#ZAP70 dephosphorylation (at membrane)
R(b).Z(Y∼4P) -> R(b).Z(Y∼3P) km
R(b).Z(Y∼3P) -> R(b).Z(Y∼2P) km
R(b).Z(Y∼2P) -> R(b).Z(Y∼P) km
R(b).Z(Y∼P) -> R(b).Z(Y∼U) km

#ZAP70 dephosphorylation (in solution)
Z(b1,b2,Y∼4P) -> Z(b1,b2,Y∼3P) kc
Z(b1,b2,Y∼3P) -> Z(b1,b2,Y∼2P) kc
Z(b1,b2,Y∼2P) -> Z(b1,b2,Y∼P) kc
Z(b1,b2,Y∼P) -> Z(b1,b2,Y∼U) kc

end reaction rules

begin actions
generate_network({overwrite=>1});
writeMfile({});

end actions
~~~

